# Kinetic model of small RNA-mediated regulation suggests that a small RNA can regulate co-transcriptionally

**DOI:** 10.1101/2020.11.03.365106

**Authors:** Matthew A. Reyer, Shriram Chennakesavalu, Emily M. Heideman, Xiangqian Ma, Magda Bujnowska, Lu Hong, Aaron R. Dinner, Carin K. Vanderpool, Jingyi Fei

## Abstract

Small RNAs (sRNAs) are important regulators of gene expression in bacteria, particularly during stress responses. Many genetically and biochemically well characterized sRNAs regulate gene expression post-transcriptionally, by affecting translation and degradation of the target mRNA after they bind to their targets through base pairing. However, how regulation at each of these levels quantitatively contributes to the overall efficacy of sRNA-mediated regulation is not well understood. Here we present a general approach combining imaging and mathematical modeling to determine kinetic parameters at different levels of sRNA-mediated gene regulation. Unexpectedly, our data reveal that certain previously characterized sRNAs are able to regulate some targets co-transcriptionally, rather strictly post-transcriptionally, and suggest that sRNA-mediated regulation can occur early in the mRNA’s lifetime, perhaps as soon as the sRNA binding site is transcribed. In addition, our data suggest several important kinetic steps that may determine the efficiency and differential regulation of multiple mRNA targets by an sRNA. Particularly, binding of sRNA to the target mRNA is likely the rate-limiting step and may dictate the regulation hierarchy observed within an sRNA regulon.

## Introduction

To cope with changes in both natural and host environments, microbes have evolved diverse mechanisms to sense, respond to, and adjust to stress conditions. Small RNAs (sRNAs) are common mediators of gene regulation in bacteria, especially in stress responses, and have been observed to provide survival benefits during infections, biofilm formation, and exposure to toxins and antibiotics (*1*–*5*). In the canonical scheme of sRNA-mediated gene regulation (Figure 1A), sRNAs, often along with a chaperone protein, Hfq, target and bind mRNAs via incomplete Watson-Crick base-pairing (*6–8*). As many sRNA binding sites on target mRNAs partially overlap with the ribosome binding site (RBS), binding of sRNAs can affect mRNA translation. In addition, the stability of the mRNAs can be affected through RNase E-mediated co-degradation of the sRNA-mRNA complex (*6*–*8*). Previous biochemical studies suggest two mechanisms for sRNA-mediated degradation: (1) sRNA-mediated reduction of translation leads to a change in degradosome access to the target mRNA, thereby increasing the degradation rate of sRNA-bound mRNA (*9*–*12*) (here referred to as “passive degradation”, or “translation-coupled degradation”, interchangeably), and (2) Modulation of degradation through direct recruitment of the degradosome (*9*,*13*–*16*) or direct obstruction of RNase E cleavage sites (*17,18*) (here referred to as “active degradation”). For a particular target mRNA, distinct sRNAs may regulate at one or more levels of expression – translation or mRNA stability – by different molecular mechanisms (*7*,*8*,*13*,*14*). However, how control at each of these levels quantitatively contributes to the overall efficacy of sRNA-mediated regulation is not well characterized.

**Figure 1.**
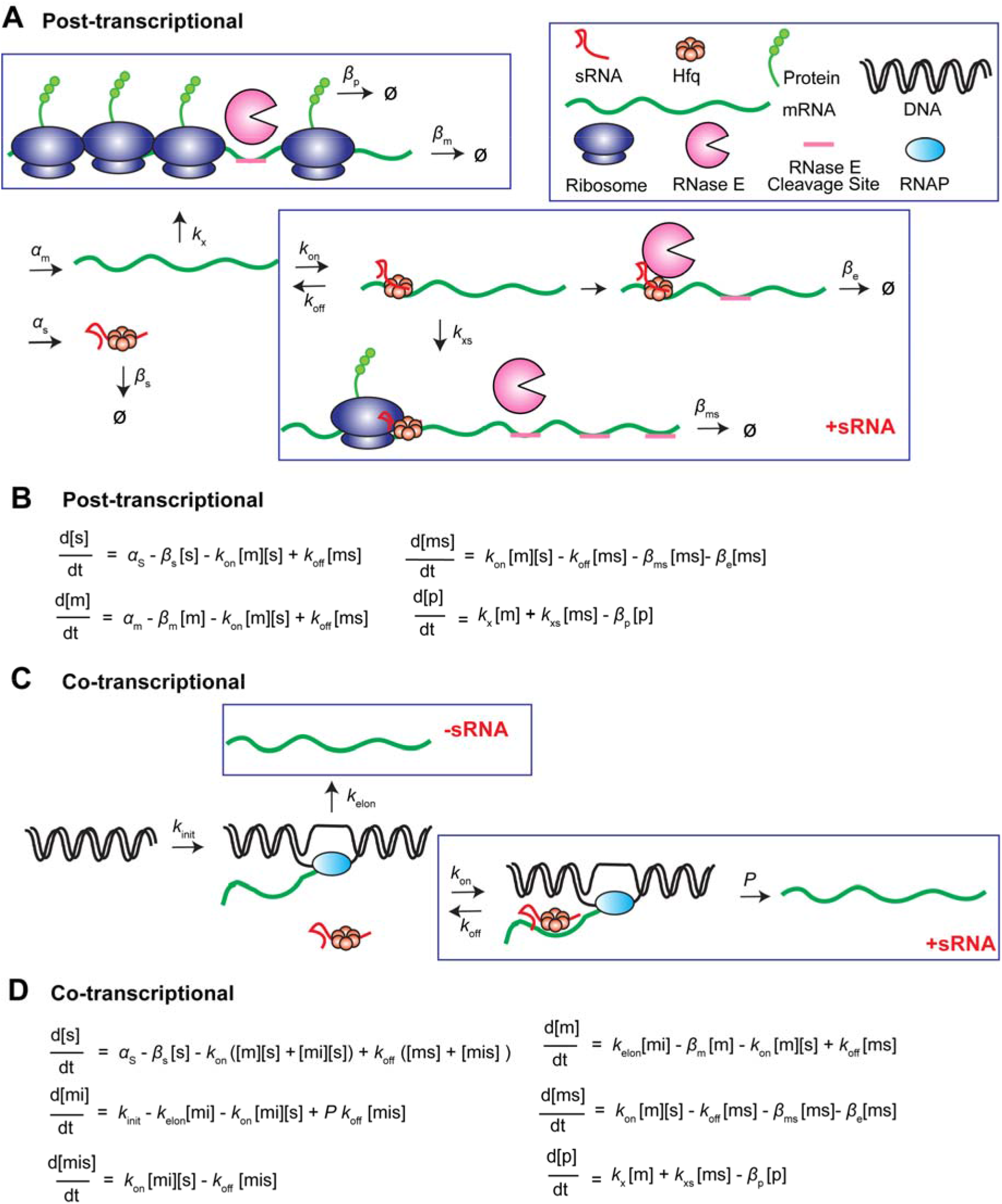
Model for determination of kinetic parameters of sRNA-mediated regulation *in vivo*. (A) Kinetic model describing sRNA-mediated, post-transcriptional regulation. (B) ODE for post-transcriptional regulation model. (C) Kinetic model for co-transcriptional regulation. (D) ODE for co-transcriptional regulation model. Parameters are described in the text.

One characteristic feature of sRNA regulators is their ability to regulate multiple target mRNAs in the same regulon (*19*–*22*). Previous studies have shown that the regulation of various targets by the same sRNA can exhibit a hierarchical pattern; *i.e*., certain targets are more effectively regulated than others (19,*23*,*24*). Such prioritization in regulation helps optimize stress responses when sRNA abundance is limited (*25*). However, the *in vivo* kinetic determinants that set the regulation hierarchy are largely unclear. A previous *in vivo* kinetic characterization of the sRNA target search and sRNA-mRNA co-degradation processes suggests that the *in vivo* binding affinity between specific sRNA-mRNA pairs can contribute to setting the regulatory hierarchy (*26*), whereas the *in vitro* binding affinity does not seem to correlate with the regulation hierarchy (*19*). In addition, a recent RIL-seq (<u>R</u>NA <u>I</u>nteraction by <u>L</u>igation and sequencing) based study found a positive correlation between the Hfq occupancy of the target mRNA and sRNA-target interaction frequency, indicating that the binding efficiency of Hfq may affect the regulation priority of the target mRNA (*27*). These observations suggest that *in vivo* target search and regulation kinetics may be collectively determined by complex molecular interactions and kinetic pathways that are difficult to fully recapitulate *in vitro* and therefore require *in vivo* characterization.

In this work, we sought to provide a comprehensive model of sRNA-mediated regulation at the level of translation and mRNA stability via different mechanisms. To achieve this goal, we utilized a genetically and biochemically well characterized *E. coli* sRNA, SgrS, as a model. SgrS is the central regulatory effector of the glucose-phosphate stress response. Intracellular accumulation of phosphorylated glycolytic intermediates, such as the phosphorylated glucose analog α-methyl glucoside-6-phosphate (αMG6P), along with depletion of other glycolytic intermediates, launches transcription of SgrS, and subsequent regulation of several mRNA targets (*28*). The best characterized targets include negatively regulated *ptsG* mRNA (encoding glucose transporter) (*14*,*29*), *manXYZ* mRNA (encoding mannose transporter) (*30–33*), *purR* mRNA (encoding purine biosynthesis operon repressor) (*34*,*35*), as well as positively regulated *yigL* mRNA (encoding a phosphatase that can dephosphorylate non-metabolizable sugars so they can be excreted to relieve stress) (*35*).

By implementing a combined single-cell imaging and mathematical modeling approach, we determined the kinetic parameters of SgrS regulation of a subset of its target mRNAs. Unexpectedly, our data reveal that SgrS is able to regulate some targets co-transcriptionally, rather than only acting on fully synthesized transcripts. Examination of another sRNA, RyhB, further suggests that acting on nascent mRNA co-transcriptionally may be a general feature for sRNAs previously believed to be post-transcriptional regulators. In addition, our data suggest several important kinetic steps that may determine the efficiency and differential regulation of multiple mRNA targets by an sRNA. Particularly, binding of sRNA to the target mRNA is likely the rate-limiting step and may dictate the regulation hierarchy observed within an sRNA regulon. Our approach may be used as a general platform for dissecting kinetic parameters and providing mechanistic details for sRNA-mediated regulation.

## Materials and methods

### Bacterial strains, plasmids

DB166 was made via P1 transduction by moving *lacIq*, *tetR, specR* cassette from JH111 (*30*) into DJ480. *ΔryhB∷cat* was moved to DB166 from EM1453 (*36*) via P1 transduction to create DB186. *rne701-FLAG-cat* was moved into strains DB166 and JH111 (*30*) by P1 transduction from TM528 (*37*) to create XM100 and XM101 respectively. The *ryhB::tet* allele in strain XM221 was created by using primers OXM211 and OXM212 with homology to RyhB to amplify the tetracycline resistance cassette. The PCR product was recombined into the chromosome of XM100 using λ red functions provided by pSIM6 (*38*).

Target mRNAs are all encoded by pSMART plasmid and under P_tet_ promoter. Target mRNA reporters carry the small RNA binding sequence from the endogenous mRNAs, and a *sfGFP* gene (Supplementary Figure S1). pSMART_*ptsG*-10aa-*sfGFP* (“10aa” refers to the first 10 codons) was generated from pZEMB8 (*19*) using site directed mutagenesis and the pSMART LCKan Blunt Cloning Kit (Lucigen, 40821-2). Briefly, the lac promoter of pZEMB8 was switched to a tet promoter to reduce leaky expression, using primers (JZ25 and JZ26) that include 5’ overhangs containing the tetracycline promoter sequence. The fragment containing the entire promoter, gene of interest, and terminator was generated by PCR using primers EH1 and EH2 and ligated into the pSMART vector, following manufacture’s instructions. pSMART_*manX*-34aa_*sfGFP* was generated following the same method as pSMART_*ptsG*-10aa_*sfGFP*, with pZEMB10 (*35*) serving as the template for the *manX*-34aa-*sfGFP* region, and primers JZ26 and EH3 containing the tetracycline promoter sequence. pSMART_*ptsG*-10aa-*sfGFP* was further used to generate pSMART_*purR*-32aa-*sfGFP* and pSMART_*sodB*_430_-*sfGFP* using Gibson Assembly. *sodB*_430_ contains RyhB binding site on *sodB* mRNA and additional 363 nucleotides in the coding region. pSMART_*sodB*_130_-*sfGFP* and pSMART_sodB_130+30_-*sfGFP* were generated from pSMART_*sodB*_430_-*sfGFP* by using primers (EH390/EH391 and EH440/441) that amplify the entire plasmid, excluding the regions that were not desired in *sodB*_130_-*sfGFP* or sodB_130+30_-*sfGFP*. The PCR products were then phosphorylated (NEB M0201S) and ligated (NEB M0202S) before transformation. Each plasmid was confirmed by DNA sequencing and transformed into the various genetic backgrounds utilized in this study.

All cell strains and plasmids used in this work are listed in Supplementary Table S1, and primers used for PCR are listed in Supplementary Table S2.

### Culture growth and induction for imaging experiments

For all imaging and qPCR experiments, overnight *E. coli* cultures were grown in LB media with 25 ug/mL Kanamycin. Overnight cultures were diluted 100 fold in MOPS-Minimal media (TEKnova, M2106) supplemented with 1% glycerol and 25 μg/mL kanamycin at 37 °C. The cells were grown to approximately OD = 0.2-0.3, at which point SgrS or RyhB was induced by adding 0.5% aMG or 500 μM DIP directly to the culture. The stress was present for 30 minutes before induction of the reporter mRNA construct using 10 ng/mL anhydrous tetracycline (aTc, Sigma-Aldrich). The time of aTc induction marked the t=0 time point in imaging experiments. Fractions of cells were taken at different time points after mRNA induction for downstream sample treatment.

### Fluorescence *in situ* hybridization (FISH)

10 FISH probes targeting the sfGFP coding region, 9 probes for SgrS and 4 probes for RyhB were designed using the Stellaris Probe Designer from Biosearch, and labeled as previously described (*26*). sfGFP probes were labeled Alexa Fluor 568 NHS ester (A568, Invitrogen A20003). SgrS and RyhB probes were labeled with Alexa Fluor 647 NHS ester (A647, Invitrogen A20006). The16S rRNA probe was labeled with Alexa Fluor 405 NHS ester (A405, Invitrogen A30000). The A405 signal serves to indicate sufficient permeabilization. FISH was performed as previously described (*26*). 10 mL of culture of cells were taken out at the corresponding time points and fixed with 4% formaldehyde at room temperature (RT) for 30 minutes. Cells were then permeabilized with 70% ethanol for 1 hour at RT. After ethanol permeabilization, 60 μL samples were taken for each time point and cells were additionally permeabilized with 25 μg/mL lysozyme for 10 minutes (1 μg/mL lysozyme corresponds to 70 units/mL). Cells were hybridized with labeled DNA probes (Supplementary Table S2) in the FISH Hybridization buffer (10% dextran sulfate (Sigma D8906) and 10% formamide in 2x SSC) at 30° C in the dark for overnight. The concentration of the labeled probes was 15 nM per probe for mRNAs, 50 nM per probe for sRNAs, and 10 nM for 16S rRNA. After the hybridization, samples were washed three times with 10% formamide in 2x SSC, and resuspended in 4x SSC.

### Epi-fluorescence Imaging and image analysis

Cells in 4x SSC buffer were imaged in 3D printed 2-well chambers. 1.2-1.4 μL of the sample were placed on the glass slide bottom of the chamber, with a 1% agarose gel pad placed on top to lay the cells flat. Imaging was performed on a custom inverted microscope (Nikon Ti-E with 100x NA 1.49 CFI HP TIRF oil immersion objective) (*39*). Multicolor Z-stack images were taken with 0.130 μm step size and 11 slices for each color. SgrS-A647 and RyhB-A647, mRNA-A568, sfGFP, and 16S rRNA-A405 were imaged with a 647 nm laser (Cobolt 06-01), a 561 nm laser (Coherent Obis LS), a 488 nm laser (Cobolt 06-01), and a 405 nm laser (CrystaLaser, DL405-025-O), respectively. In addition to the multicolor z-stack images, each image had a corresponding differential interference contrast (DIC) image, used for segmentation and image analysis purposes.

Cells were segmented individually based on DIC images using homemade MATLAB code (*40*). The segmented cell mask was then overlaid on each color channel stack individually, and the volume-integrated fluorescence intensity was calculated by adding the area-integrated intensities of each cell for the 5 most in-focus slices (the most in-focus slice, and two slices above and below). The background intensities of the image and of the cells due to nonspecific binding of the FISH probes were subtracted from the calculated volume-integrated intensities. The signal contributed by probe nonspecific binding was measured using the same imaging conditions by calculating the volume integrated intensities of cells lacking target RNAs but in the presence of the FISH probes at the same concentration as for positive samples. Δ*sgrS* cells (JH111) without transformation of any mRNA-*sfGFP* fusion plasmids were used for background measurements in the sRNA, mRNA, and GFP channels. The 16S rRNA-A405 signal was used as an indicator of sufficiently permeabilized and labeled cells. Background A405 fluorescence intensity distribution due to probe nonspecific binding was first determined using cells labeled with the same concentration of off-target A405-labeled probes. A threshold at the 90^th^ percentile of the background intensity distribution was then used as the 405 intensity cutoff. Cells with 16S rRNA-A405 intensities below this threshold (less than 10% of the total population) were considered not sufficiently permeabilized, and not included in further analysis.

### SMLM Imaging and image analysis

SMLM imaging was conducted using the same microscope as described above with super-resolution modality (*39*). Fixed cells were immobilized on the 8-well chambered glass coverslip (Cellvis C8-11.5H-N) using poly-L-lysine (Sigma-Aldrich P8920), and imaged in imaging buffer (50 mM Tris-HCl, 10% glucose,1% 2-Mercapgtoethanol (Sigma-Aldrich M6250), 50 U/mL glucose oxidase (Sigma Aldrich G2133-10KU), 404 U/mL catalase (EMD Millipore 219001) in 2X SSC, pH = 8.0). Images were acquired through a custom programmed data acquisition code, which programs the laser power, camera exposure time, and spot detection threshold, using the Nikon NIS JOBS function. SMLM images were reconstructed with the IDL analysis package as previously published (*39*).

### RT and qPCR

Total RNA was extracted from each sample using Trizol (Thermo Fisher, 15596026) extraction. 2 mL culture of bacterial cells were collected at the desired time point and immediately spun at 12,000 g for 1 minute in cold. The cell pellet was homogenized in 200 μL of trizol incubated at RT for 5 minutes. 1/5 volume of chloroform was added to the Trizol mixture. After incubation for 2-5 minutes at RT, the mixture was centrifuged at 12,000 g for 5 minutes. The upper phase was transferred to a new tube, and extracted again with chloroform. The aqueous layer was collected, from which the RNA was then precipitated by standard ethanol precipitation. The total RNA pellet is resuspended in nuclease-free water, and further desalted by a P6 microspin column (Bio-Rad, 7326221). Genomic DNA contamination in the total RNA was further removed by DNase treatment. 2 μL of Turbo DNase (Thermo Fisher, AM2238) was added to 2 μg of total RNA, and the reaction was incubated for 2 hours at 37° C. The DNase was inactivated by adding EDTA (pH = 8) at a final concentration of 15 mM, and incubating at 75° C for 10 minutes. The reaction was desalted by a P6 column.

Each reverse transcription (RT) reaction was performed using 50 ng total RNA in 1 mM dNTPs (NEB N0447S), 10% DMSO (Fisher, BP231), 10 mM DTT (Sigma-Aldrich, 10197777001), 250 nM of gene specific reverse primer (IDT), and 20-fold dilution of reverse transcriptase from iScript cDNA Synthesis Kit (Bio-Rad, 1708891) and incubated following manufacturer instructions. Each qPCR reaction was prepared using 1X SsoAdvanced Universal SYBR Green Supermix (Bio-Rad 1725274), 250 nM forward and reverse primers (Supplementary Table S2), and 1 μL of cDNA generated by the RT reaction in a final volume of 20 μL. The qPCR reactions were performed with CFX real-time PCR system (Bio-Rad), using pre-incubation of 95 °C for 30 s, followed by 40 cycles of 95 °C for 10 s and 60 °C for 30 s. The reported D/U ratio (R_D/U_), a ratio between the downstream and upstream amplification of the mRNA target, was calculated as:

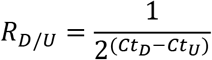

where Ct_D_ and Ct_U_ are the Ct values of the downstream and upstream amplicons respectively.

### Determination of sRNA and mRNA copy numbers

To convert the mRNA and sRNA fluorescence values to molecule copy numbers, a qPCR calibration curve of RNA copy number vs. Ct value was first built. *ptsG-sfGFP* mRNA and SgrS were produced using *in vitro* transcription. PCR using forward primers harboring the T7 promoter sequence were used to produce linear dsDNA transcription templates (Supplementary Table S2) and 1 μg template was incubated in T7 buffer (160 mM HEPES-KOH, pH 7.5, 20 mM DTT, 3 mM each rNTP, 20 mM MgCl_2_, 2 mM spermidine, 120 unites SUPERase In RNAse inhibitor) and 10 units T7 RNA polymerase (kind gift from Yuen-Ling Chan) at 37 °C for overnight. 4 units TURBO DNase was added to remove template DNA, and incubated at 37 °C for an additional 2 hours. RNA was extracted using standard phenol-chloroform and confirmed on a 7% Urea-PAGE gel.

To build a calibration curve between Ct value and RNA copy number, RT reactions were performed on a series of dilutions of *in vitro* transcribed RNA, from 10 ng to 0.001 ng. Different amounts of *in vitro* transcribed RNA were spiked into collected cell samples, then subjected to the same total RNA extraction protocol as described above. Briefly, *JH111* cells (Δ*sgrS* cells with the plasmid encoding the mRNA-*sfGFP)* were grown under the same conditions used for imaging and collected when cells reached OD_600_ = 0.2-0.3. Cells were spun down, then homogenized in Trizol. At this point (after adding Trizol, but before subsequently spinning down and adding chloroform) the *in vitro* transcribed RNA was added. RT was performed using iScript cDNA Synthesis Kit (Bio-Rad, 1708891) and qPCR was performed using SsoAdvanced Universal SYBR Green Supermix (Bio-Rad 1725274). A linear function was fit between the Ct values of the qPCR reactions and the logarithm of the input RNA copy numbers (Supplementary Figure S2A). The copy number of the RNA was calculated using the known molecular weight of the RNA and the amount of RNA added to the initial RT reaction.

To relate RNA copy number and arbitrary fluorescence values, cell samples with different RNA expression levels were subjected to RNA extraction, RT-qPCR, and fluorescence measurement, as described above. Based on the Ct value *vs.* RNA copy number calibration curve built above, *sfGFP* fusion mRNA and SgrS copy numbers were calculated for the extracted RNA of each sample, and further converted into copy number per cell based on the cell numbers measured by OD_600_ for each sample. RNA copy number per cell was then plotted against the volume-integrated cell fluorescent intensities for each corresponding sample and fit with a linear function (Supplementary Figure S2B). Fluorescent intensities of the cells from the imaging experiments were compared to this calibration curve of fluorescent intensity *vs*. RNA copy number to extract RNA copy number per cell.

### Simulation, fitting, and model selection

We used Markov Chain Monte Carlo (MCMC) simulation to explore the parameter spaces of our kinetic models as defined by their ordinary differential equations (ODEs). Specifically, we utilized the *emcee* package (*41*), which is a Python implementation of the Goodman-Weare Affine Invariant Ensemble Sampler (*42*), and integrated the ODEs with the LSODA solver (*43*,*44*). In this approach, an ensemble of parameter sets evolves to sample a Bayesian posterior distribution, which is the product of a prior distribution and a likelihood function. Assuming Gaussian and independent errors, the logarithm of the likelihood (log-likelihood) function takes the form:

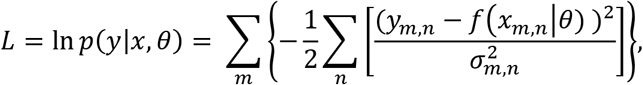

where *m* is the molecular species (mRNA, sRNA, and protein in the WT and *rne*701 strain, for a total of 6), *n* is the time point (7 in our case, *t* = 0, 1, 3, 6, 12, 18 and 24 min), *y*_m,n_ is the experimental value for molecular species *m* at time t_n_, (in units of copy number for sRNA and mRNA, and arbitrary fluorescent unit for protein), *f*(*x_m,n_*|*θ*) is the simulated value for molecular species *m* at time *t_n_* given the parameter value set *θ*, 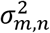 is the experimental variance for molecular species m at time point *t_n_*. The log-posterior distribution is the sum of the log-prior distribution and log-likelihood function.

We fit parameters by running simulations in a two-step process. First, mRNA transcription and translation rates were fit using the –sRNA experimental data, *i.e.* the data acquired from the cell samples in the absence of sRNA. The best fit parameter values and their associated errors were used as prior distributions for transcription and translation rates in the second step, where the rest of the parameters were determined by fitting to the +sRNA experimental data, acquired from cell samples in the presence of sRNA. For the co-transcriptional regulation model using the one-step transcription module, the –sRNA simulations explored a 3-dimensional parameter space: [*α*_m_, *k*_x_, *β*_m_]; and the +sRNA simulations explored a 9-dimensional parameter space: [*k*_on_, *k*_off_, *k*_xr_, *β*_e_, *β*_ms_, *k*_x_wt_, *k*_x_rne_, *α*_ms_wt_, *α*_ms_rne_], where *k*_xr_ is the ratio *k*_xs_/*k*_x_. For the post-transcriptional regulation model using the one-step transcription module, *α*_ms_wt_ and *α*_ms_rne_ were set to *α*_m_wt_ and *α*_m_rne_, respectively. For the co-transcriptional regulation model using the two-step transcription module, the –sRNA simulations explored a 3-dimensional space: [*k*_init_, *k*_x_, *β*_m_]. The elongation rate, *k_elon_* was assumed to be a constant for each mRNA, determined by dividing a constant elongation speed (50 nucleotides per second (*45*)) by the length of the mRNA. The +sRNA simulations explored a 9-dimensional space [*k*_on_, *k*_off_, *k*_xr_, *β*_e_, *β*_ms_, *k*_x_wt_, *k*_x_rne_, *P*_wt_*, P*_rne_], where *P*_wt_ and *P*_rne_ represent the probability of generating full length mRNA in WT *rne* and *rne701* backgrounds, respectively. For the post-transcriptional regulation model using the two-step transcription module, *P*_*wt*_ and *P*_*rne*_ were set to 1. For –sRNA simulations in 3 dimensions, 50 walkers, representing 50 parameter sets, each evolved for 10000 steps, which we found to be a sufficient number of steps for the log posterior to level off. For +sRNA simulations in 9 dimensions, 100 walkers each evolved for 10000 steps. Initial positions for the walkers were chosen at random from the bounded interval of possible values defined by its prior distribution. We used the default settings for the *emcee* sampler, such that the each move is a “stretch” move, with stretch parameter, *a* = 2, giving an average acceptance fraction equal to 0.44 (*41*,*42*).

For –sRNA fitting, the prior distributions for the free parameters were uniform distributions (Table S4). For the +sRNA fitting, the prior distributions of the parameters determined from the – sRNA fitting were normal distributions centered on their –sRNA maximum a posteriori (MAP) values, and the prior distributions for the remaining parameters were uniform distributions. After the parameter fitting, the posterior probability distributions of the fitted parameters were determined, along with their MAP values and associated errors. For experimentally determined variables, the widths of the normal distributions were determined by their experimental errors. For the remaining free parameters, the widths of the uniform distributions were set empirically, either by observing physical constraints (e.g., *k*_*on*_ is constrained by the diffusion limit) or by logical constraints (e.g., *k*_*xr*_ cannot be below 0 or above *k*_*x*_).

Each experimental replicate was fit separately. 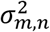 was the same across all replicates. A single set of parameter values was chosen to be the best fit for the combined samples by selecting the point estimate of the MAP parameter values for the best walker for each replicate, then averaging over the replicates. One replicate in a –sRNA simulation was one experimental dataset containing mRNA and associated protein values, with datasets for WT and *rne*701 backgrounds fit separately. One replicate in a +sRNA simulation was a combination of one experimental dataset in the WT background, and one in the *rne*701 background. The reported parameter values and their associated errors were the mean and standard deviations of the MAP values from all simulations, respectively. All simulations were performed with custom software written in Python, and parallelization was implemented using *emcee*. We utilize both CPU and GPU functions to maximize the efficiency of our simulations. All codes for all simulations are available publicly on GitHub: (https://github.com/JingyiFeiLab/Regulation_Kinetics).

The Bayesian information criterion (BIC) was used for model selection between post-transcriptional and co-transcriptional regulation models. The BIC is defined as:

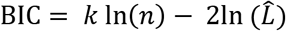

where 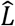 is the maximized likelihood value of the model, *k* is the number of parameters (*k* = 7 for post-transcriptional model, *k* = 9 for co-transcriptional model, accounting for the added variables *P*_*wt*_ and *P*_*rne*_), *n* is the number of data points or observations (*n* = 42 in our case, representing 7 time points × 3 molecules × 2 *rne* backgrounds). For each target, the minimized BIC was calculated for both the post- and co-transcriptional models, and the model which produced the lowest BIC was selected.

## Results

### Kinetic model and experimental measurement of sRNA-mediated regulation

Since SgrS has been biochemically characterized as a post-transcriptional gene regulator, we first setup a post-transcriptional regulation model to describe this process, including regulation at the levels of both translation and degradation (Figure 1A). In the absence of the sRNA, this model includes basal levels of mRNA transcription (*α*_m_, as rate constant), translation (*k*_x_), and endogenous mRNA and protein degradation (*β*_m_ and *β*_p_ respectively). When the sRNA is produced, its transcription rate is defined by *α*_s_ and the effective degradation rate by *β*_s_. *β*_s_ approximates endogenous degradation as well as target-coupled degradation with all other mRNA targets except for the specific target mRNA of interest. The sRNA binds to an mRNA target with an on-rate of *k*_on_ and dissociates with an off-rate of *k*_off_. Upon binding, the translation activity of the bound mRNA changes to *k*_xs_. The sRNA-mediated degradation is described by *β*_ms_ for translation-coupled degradation and *β*_e_ for active degradation.

Production of the sRNA, SgrS (from the endogenous chromosomal gene), was induced by exposing *E. coli* cells to glucose-phosphate stress using α-methylglucoside (αMG) (*14*). The target mRNAs (containing SgrS binding sequences) fused to the super-folder (sf) GFP gene (*46*) were carried on low-copy number plasmids under the control of a tetracycline promoter (P_*tet*_) (Figure 2A and Supplementary Figure S1). In contrast to the induction scheme commonly used in previous studies in which the changes in target mRNA abundance or translation were recorded after sRNA induction, we chose to induce SgrS before target mRNA induction and then record the levels of SgrS, target mRNA and protein simultaneously as a function of time. Time t = 0 was defined as the time at which the target mRNA was induced (Figure 2B). Fractions of cells were fixed at different time points. SgrS and the target mRNAs were fluorescently labeled with DNA oligo probes through a standard fluorescence in situ hybridization (FISH) method (*26,47*). Translation of the *sfGFP* fusion produced a direct fluorescent readout for protein levels (Figure 2B). The single-cell sRNA, mRNA, and protein levels were characterized by their volume-integrated fluorescent signals (*40*). sRNA and mRNA copy numbers were further determined by comparing fluorescent intensities and the Ct values in the reverse transcription and quantitative PCR (RT-qPCR)-based calibration (Figure 2C and Supplementary Figure S2). As the copy numbers of sRNA and mRNA were in the range of tens to hundreds per cell, we described the time-dependent changes in sRNA, mRNA and protein deterministically by mass action equations (Figure 1B).

**Figure 2.**
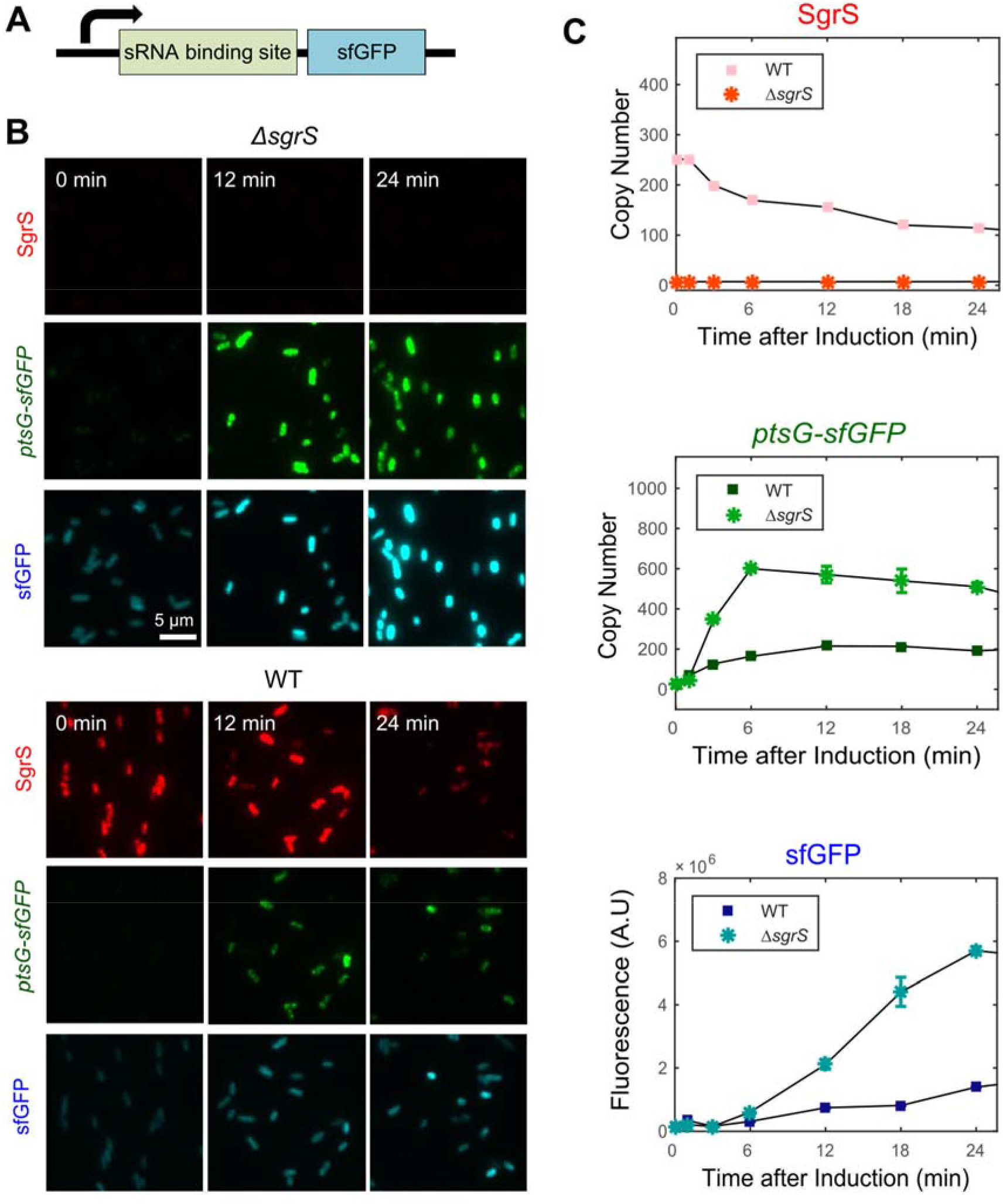
Illustration of experimental setup and representative results. (A) Illustration of the target mRNA, including the 5’ UTR and part of the coding region from the endogenous mRNA target containing the SgrS binding site, and a coding region for sfGFP reporter. (B) Representative images of SgrS (red), *ptsG-sfGFP* mRNA (green) and sfGFP signal (blue) in the absence (upper) or presence (lower) of sRNA induction over 24 minutes. (C) Measured sRNA, mRNA, and protein levels from images in (B), representing volume-integrated single cell fluorescence values, converted to copy numbers for the case of RNA molecules.

We chose to pre-induce the sRNA for two reasons. First, by capturing the sRNA-mediated changes in the production of new proteins, we can more accurately measure regulation at the translational level. sRNA-mediated regulation generally occurs within minutes (*14*,*24*,*48*). However, many proteins, including the reporter *sfGFP*, have long lifetimes *in E.coli*, which are essentially determined by rate of dilution due to cell division (*49*). Therefore, the fluorescent signal from already existing proteins in the cell can overwhelm any protein level changes caused by sRNAs. More importantly, we were interested in the timing of sRNA-mediated regulation of target mRNAs and more specifically, whether sRNA can act on the newly synthesized mRNAs co-transcriptionally. In the case of pre-induced mRNA, the mature mRNAs outcompete the nascent mRNAs owing to their relative abundances, which may make any effect at the transcriptional level undetectable.

For each sRNA-mRNA pair, we measured the time-course changes of sRNA, mRNA and protein levels in four genetic backgrounds: wild-type (WT), Δ*sgrS*, *rne*701, and *rne*701 Δ*sgrS*. Time-dependent changes in mRNA and protein upon mRNA induction were recorded in the absence of SgrS for the determination of parameters describing basal transcription (*α*_m_) and translation (*k*_x_) activities. By comparing the fusion mRNA and protein levels in the Δ*sgrS* strain in the presence of αMG with the corresponding levels in the WT strain in the absence of αMG, we noticed that the presence of αMG alone (*i.e.*, without ensuing production of SgrS) reduced the efficiency of induction of the mRNA fusion (Figure 2C, Supplementary Figure S3, and Supplementary Table S3). Therefore, to quantify the regulation by the sRNA specifically, we use the Δ*sgrS* and *rne*701 Δ*sgrS* grown in the presence of αMG as our “-sRNA” condition to quantify the basal transcription and translation activities of the target mRNA in the WT and *rne*701 backgrounds, respectively. Comparison of the kinetic behaviors in the WT vs. *rne*701 strain allowed us to separate the effect of sRNA-mediated passive and active degradation. The *rne*701 allele encodes a truncated RNase E protein lacking part of the C-terminal unstructured region (*50*), including RhlB, enolase, PNPase and Hfq binding sites (*16*,*51*–*53*). The *rne*701 mutant still fully retains its catalytic function, but has an impaired ability to interact with Hfq and other degradosome components without the C-terminal unstructured region. We therefore assumed that SgrS-mediated mRNA degradation is primarily through passive degradation in the *rne*701 background as previously reported (*15*,*26*,*37*).

Finally, to further constrain our model, we experimentally measured a subset of parameters. Specifically, we measured *β*_m_ using rifampicin pulse-chase experiments (Supplementary Figure S4), and *β*_s_ by first inducing SgrS and then washing away the inducer (Supplementary Figure S5). *β*_m_ of *ptsG-sfGFP* fusion mRNA did not show significant difference between the WT and *rne* mutant background, consistent with a previous finding that *rne*701 has a minor effect on endogenous mRNA degradation (Supplementary Figure S4) (*26*,*54*). *β*_s_ was slightly slower in the *rne*701 background (Supplementary Figure S5), suggesting that active co-degradation with target mRNAs contributes to the ensemble sRNA turnover, consistent with previous results (*26*). We approximated sfGFP protein half-life using the cell doubling time (~90 min) under our experimental condition (Supplementary Figure S6). *α*_s_ was determined by measuring the time-dependent production of SgrS upon induction.

### Simulation predicts that SgrS may regulate *ptsG* co-transcriptionally

Under the assumption that SgrS regulates *ptsG-sfGFP* mRNA post-transcriptionally, we fixed the *α*_m_ and *k*_x_ values obtained from the Δ*sgrS* and *rne*701 Δ*sgrS* strains and fit the rest of the parameters in the time-dependent levels of SgrS, target mRNA, and sfGFP in the WT and *rne*701 strain in the presence of αMG, including *β*_e_, *β*_ms_, *k*_xs_, *k*_on_, and *k*_off_, using Bayesian Markov Chain Monte Carlo (MCMC) modeling (Materials and Methods). However, the optimized parameters of the post-transcriptional regulation model did not accurately describe the experimental data, specifically the amplitude of sRNA-induced repression (Supplementary Figure S7A). We therefore considered an alternative model that included the possibility that SgrS could regulate its targets co-transcriptionally, rather than exclusively post-transcriptionally.

Initially, we modeled co-transcriptional regulation by allowing *α*_m_ to change in the presence of sRNA (denoted *α*_ms_). This model fit the data well (Supplementary Figure S7B). The resulting *α*_ms_ (0.87 ± 0.05 molecules•s^−1^) was smaller than α_m_ (1.9 ± 0.3 molecules•s^−1^), *i.e.*, transcription was slower in the presence of the sRNA. Since the FISH probes for the target mRNA specifically bind to the sfGFP coding region in the mRNA fusion downstream of the sRNA binding site, we infer that the reduction in the transcription rate is an indicator of sRNA-mediated regulation occurring during transcription, since only fully synthesized mRNA produce fluorescent signal. In addition, the reduction in *α*_m_, was more pronounced in the WT *rne* background (*α*_ms_ = 0.46 *α*_m_) compared to in the *rne*701 background (*α*_ms_ = 0.98 *α*_m_), suggesting that a fully assembled degradosome contributes to the strength of co-transcriptional regulation.

### SgrS decreases the abundance ratio of downstream to upstream regions relative to the SgrS binding site on the target mRNA

Since, according to our model, co-transcriptional regulation by SgrS reduces the production of full-length *ptsG-sfGFP* mRNA, we reasoned that this may be reflected by a decrease in the abundance of downstream (from the SgrS binding site) relative to upstream regions on the *ptsG-sfGFP* mRNA (henceforth referred to as the “D/U ratio”). To experimentally measure the D/U ratio, we devised a RT-qPCR assay. In this assay, we utilized two sets of primers: one amplifying the region upstream of the SgrS binding site, and the other amplifying the downstream region, in the coding region (Figure 3A). To evaluate the D/U ratio change specifically contributed by the co-transcriptional regulation, we compared RT-qPCR results on extracted RNA from cells at 1 and 15 min after induction (Figure 3B). These times were chosen based on the fact that the lifetime of *ptsG-sfGFP* mRNA is around 7-8 min (Supplementary Figure S4 and Supplementary Table S3). At 1 min after induction (D/U_t=1_), the contribution by endogenous degradation on the read-though ratio should not dominate. In addition, since the cellular level of mRNA at 1 min post-induction is low (Figure 2C), the fraction of nascent mRNAs, i.e., the mRNAs still being transcribed, compared to fully synthesized mRNAs, should be relatively high. We therefore considered this pool of *ptsG-sfGFP* mRNAs as relatively enriched in the nascent mRNAs, and expected that effects at the co-transcriptional level would be enhanced in this sample. At 15 min after induction (D/U_t=15_), *ptsG-sfGFP* mRNA levels reach steady-state, with a high cellular abundance (Figure 2C); thus the fraction of nascent mRNAs should be minimal compared to fully synthesized mRNAs. We therefore reasoned that effects at the co-transcriptional level are largely buried by effects at the post-transcriptional level.

**Figure 3.**
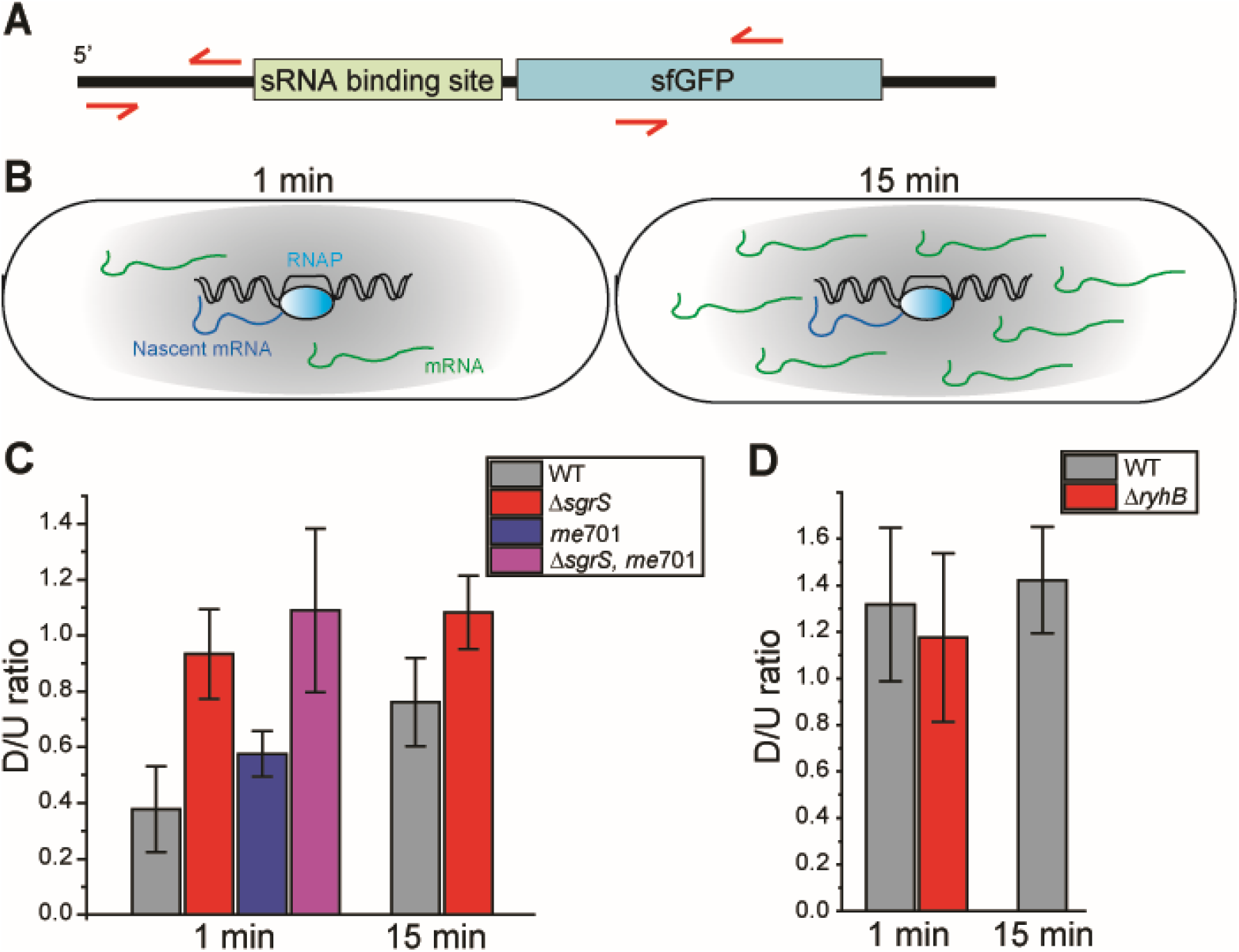
RT-qPCR measurement of the D/U ratio. (A) Schematic illustration of the qPCR primer binding sites relative to SgrS binding site on the mRNA. (B) Schematic illustration of total RNAs extracted at different time points of mRNA induction, which contain different ratios of nascent mRNAs to fully transcribed mRNAs. (C) D/U ratio in the absence and presence of SgrS. (D) D/U ratio in the absence and presence of RyhB.

We measured D/U_t=1_ and D/U_t=15_ in the WT and Δ*sgrS* cells in the presence of αMG (Figure 3C). D/U_t=15_ in the WT cell was about ~70% of that in the Δ*sgrS* cell, suggesting that the regulation by SgrS caused reduction in the abundance of the downstream region compared to the upstream region of the SgrS binding site on the target mRNA. The reduced D/U_t=15_ upon SgrS regulation may be explained by the directionality of RNase E activity, i.e., an enhanced RNase E activity on the downstream fragment with 5’ monophosphate(*55*–*57*). In comparison, D/U_t=1_ in the WT cell was about ~40% of that in the Δ*sgrS* cell, suggesting that in the nascent-mRNA enriched pool, the regulation by SgrS led to more reduction in the abundance of the downstream region compared to the upstream region, and supporting our prediction that SgrS repressed the generation of the downstream portion co-transcriptionally. As a control, D/U_t=1_ and D/U_t=15_ remained constant when inducing a non-matching sRNA, RyhB, a small RNA that is repressed by Fur (ferric uptake regulator) and produced in response to iron depletion, by adding 2,2’-dipyridyl (referred to as “DIP”) into the culture (*58*) In addition, the reduction in D/U_t=1_ was less in the *rne*701 background (~51% comparing D/U_t=1_ in the *rne*701 and *rne*701Δ*sgrS*), consistent with the predicted trend from the simulation that the co-transcriptional regulation is stronger in the WT background.

### A revised kinetic model containing co-transcriptional regulation module

After experimentally confirming the feasibility of co-transcriptional regulation, we then improved the kinetic model by linking sRNA binding to the co-transcriptional regulation (Figure 1C and D). In this revised model, we assumed that sRNA binds nascent and mature mRNAs with the same *k*_on_ and *k*_off_ rates. In order to allow for co-transcriptional binding, mRNA transcription is separated into two steps: initiation (*k*_ini_) and elongation (*k*_elon_). When nascent mRNAs are bound by sRNA during elongation, a free parameter (*P*) is introduced to the model, representing the probability of generating the full-length, mature mRNA. We allow *P* to differ between the WT and *rne*701 backgrounds. The revised kinetic model significantly improved the fitting of data for the SgrS regulation over *ptsG* (Figure 4A and B). To further validate the improved performance of the co-transcriptional regulation model, we applied the Bayesian information criterion (BIC), where a penalty is applied to the co-transcriptional model for its two added parameters (namely, *P*, in WT *rne* and *rne*701 background) (*59*), to select between co-transcriptional and post-transcriptional regulation models. The co-transcriptional model was selected by virtue of having the lower BIC value. Consistent with the qPCR results, *P* was lower in the WT background than in the *rne*701 background (Figure 4A and B, Table 1).

**Figure 4.**
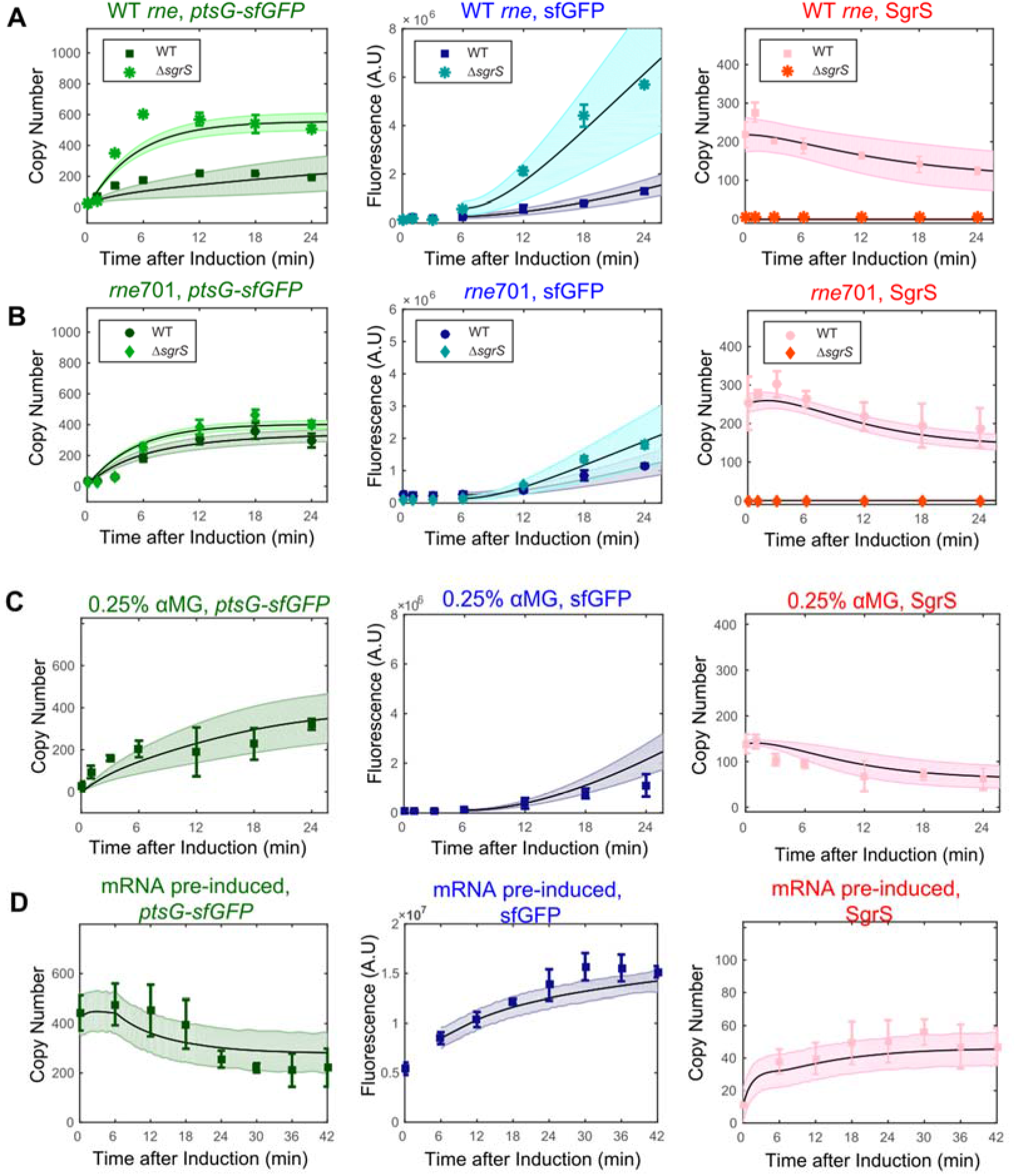
Fitting of SgrS regulation of *ptsG* expression with co-transcriptional regulation model. Time-dependent changes of SgrS, *ptsG-sfGFP* mRNA, and sfGFP levels in the presence or absence of SgrS, in the (A) WT *rne* background and (B) *rne*701 background. Points with error bars represent experimental data from 2-3 biological replicates. Each biological replicate contains ~500-1000 cells. Black lines represent fitting with the best set of parameters using co-transcriptional regulation model. Shaded, colored regions represent predicted error of the fitting, calculated by sampling from the means and errors of individual kinetic parameters. (C) Simulated prediction (black curve with shaded, colored region) using co-transcriptional regulation model for validation dataset with reduced αMG concentration for SgrS induction, overlaid with experimental data (points with error bars). (D) Simulated prediction using co-transcriptional regulation model and experimental data for validation dataset of pre-induced mRNA.

**Table 1.** parameter comparison between target mRNAs^a^.

| mRNA | *pstG* | *manX* | *purR* <sup>b</sup> | *sodB*<sub>130+30</sub> | *sodB*<sub>130</sub> |
| --- | --- | --- | --- | --- | --- |
| sRNA | SgrS | SgrS | SgrS | RyhB | RyhB |
| Repression% | 0.78 ± 0.07 | 0.53 ± 0.18 | 0.20 ± 0.05 | 0.68 ± 0.14 | 0.63 ± 0.03 |
| Steady State Protein Repression % | 0.57 | 0.43 | 0.045 | 0.67 | 0.48 |
| Steady State mRNA Repression % | 0.48 | 0.33 | 0.11 | 0.67 | 0.37 |
| $k_{on}$ (M<sup>-1</sup>s<sup>-1</sup>) <sup>c</sup> | (1.3±0.6)x10<sup>6</sup> | (3.4±1.2)x10<sup>5</sup> | (3.3±1.2)x10<sup>4</sup> | (14 ±0.6)x10<sup>5</sup> | (4.6±0.9)x10<sup>5</sup> |
| $k_{off}$ (s<sup>-1</sup>) | 0.30 ± 0.04 | 0.21 ± 0.12 | 0.56 ± 0.14 | 9.6 ± 0.3 | 9.1 ± 0.4 |
| $k_{xs}/k_x$ | 0.14 ± 0.06 | 0.28 ± 0.12 | 0.63 ± 0.13 | 0.36 ± 0.16 | 0.43 ± 0.29 |
| $\beta_{ms}$ (s<sup>-1</sup>) | (5.0±1.5)x10<sup>-3</sup> | (3.7±0.3)x10<sup>-3</sup> | (2.6±0.8)x10<sup>-2</sup> | (5.6±0.1)x10<sup>-3</sup> | (5.8±0.4)x10<sup>-3</sup> |
| $\beta_e$ (s<sup>-1</sup>) | (1.3±1.1)x10<sup>-3</sup> | (2.3±1.2)x10<sup>-3</sup> | (5.7±1.2)x10<sup>-1</sup> | (2.2±1.5)x10<sup>-3</sup> | (4.9±0.8)x10<sup>-4</sup> |
| *P* (WT) | 0.32 ± 0.14 | 0.43 ± 0.21 | N/A (Post) | 0.04 ± 0.02 | 0.16 ± 0.07 |
| *P* (*rne701*) | 0.93 ± 0.04 | 0.57 ± 0.09 | N/A (Post) | 0.80 ± 0.02 | 0.50 ± 0.23 |
| *BIC* (*co*) | 497.420 | 481.706 | 436.404 | 492.869 | 494.216 |
| *BIC* (*post*) | 577.487 | 486.493 | 429.181 | 669.051 | 515.328 |

- Reported parameter values and their associated errors are the mean and standard deviations of the outputted MAP values from all replicates. One replicate in a +sRNA simulation is a combination of one experimental dataset in WT background, and one in *rne701* background - Best fit parameters for *purR* were determined from experiments using 1% αMG, rather than the 0.5% used for other SgrS targets. This is due to the fact that repression was minor for *purR* at 0.5% αMG. The repression % for *purR* was determined using 0.5% αMG induction for a fair comparison with other SgrS targets. - $k_{on}$ is reported as M<sup>-1</sup>S<sup>-1</sup> assuming 1 molecule corresponds to 1 nM in *E. coli* cells (81).

To validate the co-transcriptional regulation model, we generated two data sets. In the first, we reduced the induction of SgrS using a lower concentration of αMG and measured *α*_s_ experimentally. In the second, we reversed the induction order of SgrS and *ptsG-sfGFP* mRNA, presenting the condition under which newly induced sRNAs regulate pre-existing mRNA targets. We simulated the time courses of SgrS, *ptsG-sfGFP* mRNA, and sfGFP using the best set of parameters obtained from a model with (Figure 4A and B) or without (Supplementary Figure S8A) co-transcriptional regulation, respectively. In both cases, the co-transcriptional regulation model predicted the experimental data better (Figure 4C and D, Supplementary Figure S8B and C).

### Co-transcriptional regulation may be a widespread mechanism utilized by sRNAs

We next asked if co-transcriptional regulation might be a general feature shared by other previously characterized post-transcriptional sRNA regulators. We applied the same imaging and modeling scheme to RyhB and one of its targets, *sodB*. We generated two fusion mRNAs, *sodB*_130_ and *sodB*_130+30_, containing the RyhB binding site and sfGFP gene (Supplementary Figure S1). *sodB*_130+30_ contains an additional 30 nucleotides which include a RNase E cleavage site and is more sensitive to RyhB regulation at the degradation level (*9*).

The responses of *sodB*_130_ and *sodB*_130+30_ to RyhB regulation were again best captured by the co-transcriptional regulation model as suggested by BIC (Figure 5A and B, Supplementary Figure S9B and C), suggesting that co-transcriptional regulation may be a general mechanism of sRNA-mediated regulation. Consistent with the SgrS regulation, co-transcriptional regulation for RyhB was also more efficient in the WT background compared to *rne*701. In addition, the *β*_e_ value of the *sodB*_130+30_ was ~4.5 fold higher than that of the *sodB*_130_, in line with the addition of the RNase E cleavage site in *sodB*_130+30_, serving as a validation of our model.

**Figure 5.**
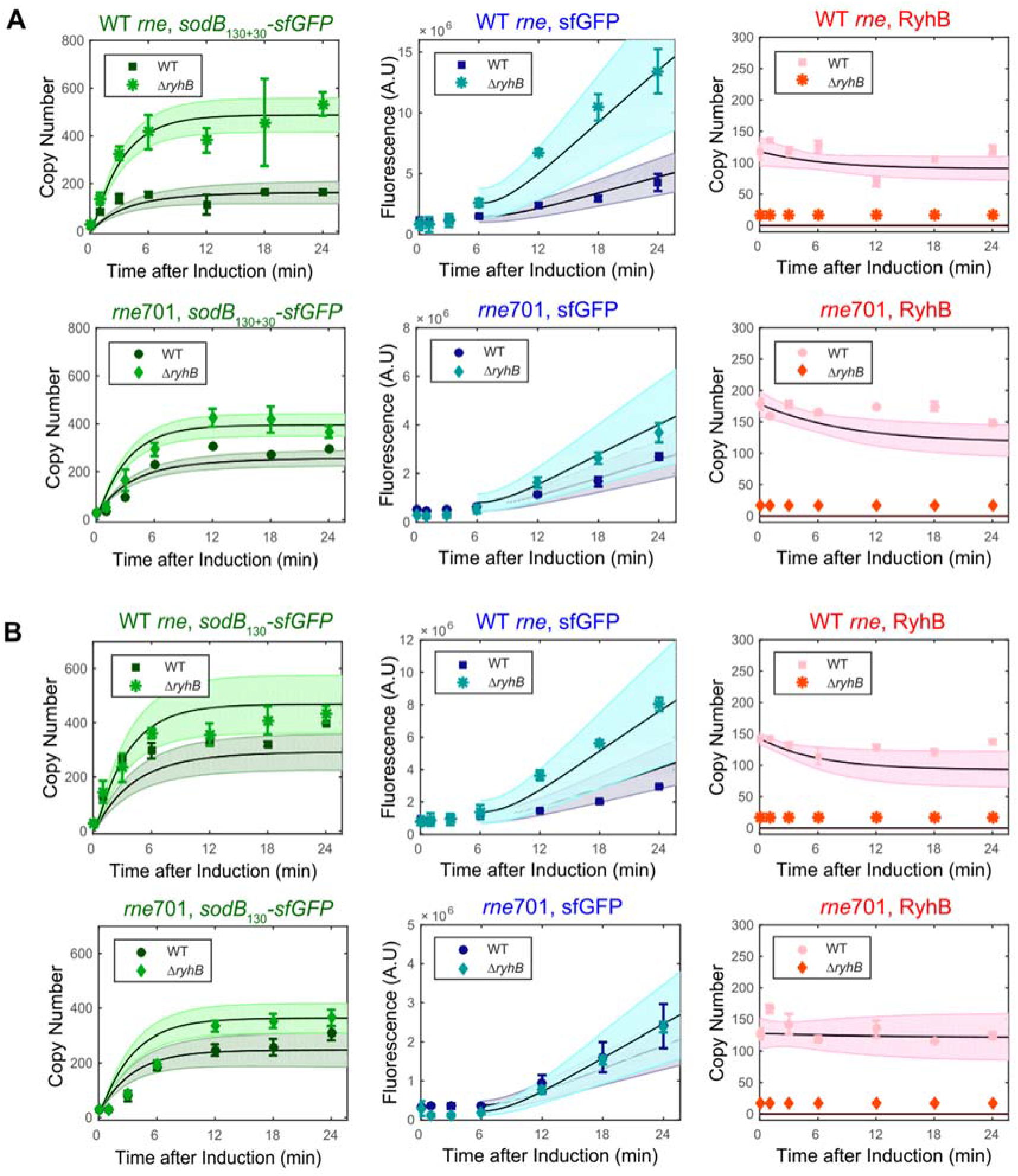
Fitting of sRNA-mediated regulation of *sodB* with co-transcriptional regulation model. (A) Time-dependent changes of RyhB, *sodB*_130+30_*-sfGFP* mRNA, and sfGFP levels in the presence or absence of RyhB, in WT *rne* (upper) and *rne*701 (lower) background. Figures follow the same format as in Figure 4. (B) Time-dependent changes of RyhB, *sodB*_130_*-sfGFP* mRNA, and sfGFP levels in the presence or absence of RyhB, in WT (upper) and *rne*701 (lower) background. Error bars represent experimental data from 2-3 biological replicates. Each biological replicate contains ~500-1000 cells.

### Parameters that contribute to regulation efficiency of sRNA over different targets

We next fit the models to two other SgrS targets, *manX* and *purR*. It has been established that *ptsG* is the primary target of SgrS, *manX* is a secondary target, and *purR* is a lower-priority target (*19*). Consistently, we observed 87%, 53% and 18% repression respectively for *ptsG*, *manX* and *purR* at the protein level at 24 minutes under the same SgrS induction condition (Table 1). At steady state, the model predicted the regulation efficiency to be 57%, 43% and 5% at the protein level, and 48%, 33% and 11% at the mRNA level for *ptsG*, *manX* and *purR,* respectively. BIC suggested the co-transcriptional regulation model better fit *manX* (Figure 6A and Supplementary Figure S9A), but the post-transcriptional model better fit *purR* (Figure 6B), indicating that the contribution of co-transcriptional regulation for *purR* is negligible.

**Figure 6.**
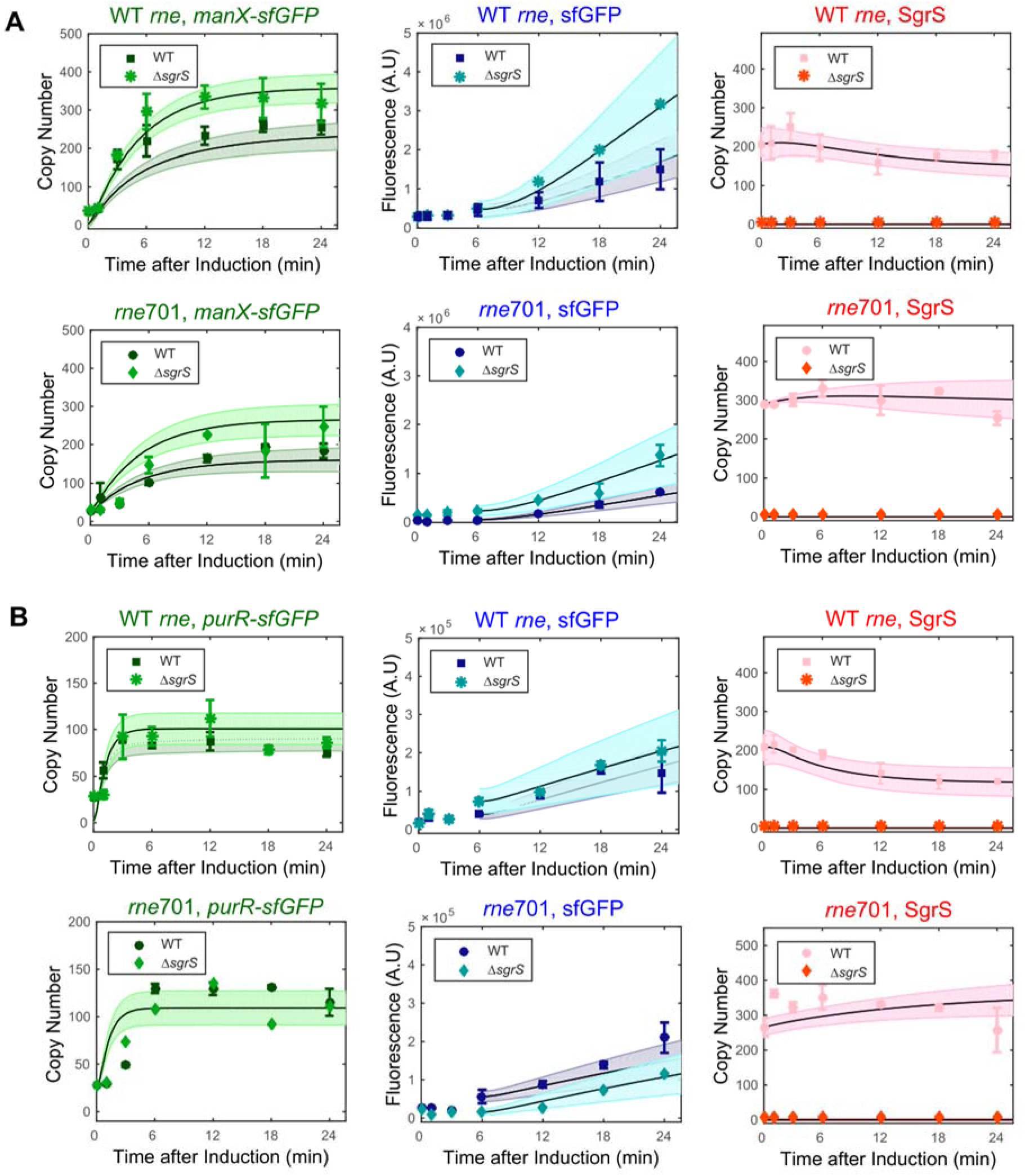
Fitting of sRNA-mediated regulation of *manX* and *purR*. (A) Time-dependent changes of SgrS, *manX-sfGFP* mRNA, and sfGFP protein levels in the presence or absence of SgrS, in WT *rne* (upper) and *rne*701 (lower) background. Figures follow the same format as in Figure 4. Regulation on *manX-sfGFP* was best fit with co-transcriptional regulation model. (B) Time-dependent changes of SgrS, *purR-sfGFP* mRNA, and sfGFP protein levels in the presence or absence of SgrS, in WT *rne* (upper) and *rne*701 (lower) background. Regulation on *purR-sfGFP* was best fit with post-transcriptional regulation model. Error bars represent experimental data from 2-3 biological replicates. Each biological replicate contains ~500-1000 cells.

Comparison of the parameters for the three mRNA targets for SgrS and two targets for RyhB (described above) suggests features that contribute to the overall regulation efficiency (Table 1).

1. Within the same sRNA regulon, a faster binding rate led to more efficient regulation. We found that *k*_on_ changed more dramatically than *k*_off_ among different targets. For SgrS, the difference in *k*_off_ was within ~2 fold among the three targets, whereas the change in the *k*_on_ values was up to ~40 fold between *ptsG* and *purR*, suggesting that the binding kinetics is dominated by *k*_on_. Interestingly, although *sodB*_130_ and *sodB*_130+30_ had the same RyhB target site, which led to the same *k*_off_, *sodB*_130+30_ showed a higher *k*_on_ than *sodB*_130_ (see Discussion). In addition, RyhB had a much higher *k*_off_ for the *sodB* constructs compared to SgrS.
2. The repression at the translation level (*k*_xs_/*k*_x_) contributed positively to the regulation efficiency among the SgrS targets. The SgrS binding site is located at the 5’ UTR on *ptsG*, partially overlapping the RBS, and within the CDS of *manX* and *purR* (*32*,*35*). SgrS inhibits translation initiation on these mRNAs through different mechanisms. On *ptsG*, base pairing of SgrS directly blocks ribosome binding, while on *manX* and *purR*, binding of SgrS guides Hfq to bind at a site close to the RBS to block ribosome binding (*32*,*35*). Our results indicate that direct binding of SgrS at the RBS may be slightly more efficient in repressing translation. *k*_xs_/*k*_x_ was similar among the two *sodB* constructs upon RyhB regulation, consistent with the fact that they share the same RyhB binding site. However, even though RyhB also regulates *sodB* through directly blocking ribosome binding at the RBS (*60*,*61*), the repression at translation level was less efficient than SgrS regulation on *ptsG*, suggesting that the different structures of formed sRNA-mRNA duplexes may affect translation to different extents.
3. For all target mRNAs, *β*_ms_ was larger than the corresponding *β*_m_, supporting the translation-coupled degradation model in which reduced translation activity upon sRNA binding leads to faster degradation of the sRNA-bound mRNA. For the three SgrS targets, there was no correlation between *β*_e_ and regulation efficiency. Although a higher *β*_e_ was observed for *purR*, the least regulated target of SgrS, a much smaller *k*_on_ value for *purR* limited the regulation efficiency. The impact of active degradation became more evident when comparing the two RyhB targets, in which most other parameters were similar. The higher *β*_e_ value of the *sodB*_130+30_ contributed to a higher regulation efficiency of *sodB*_130+30_ (67% and 67% at protein and mRNA levels for *sodB*_130+30_ respectively compared to 48% and 37% for *sodB*_130_).
4. We observed a positive correlation between the strength of co-transcriptional regulation and the overall regulation efficiency. Co-transcriptionally bound *ptsG* had a lower probability of generating a full-length mRNA compared to *manX*, while *purR* was insignificantly affected by co-transcriptional regulation. Similarly, co-transcriptionally bound *sodB*_130+30_ had a lower probability of generating a full-length mRNA compared to *sodB*_130_.

## Discussion

We have presented a general approach combining imaging and modeling, which can be used to quantify the kinetic parameters underlying differential regulation of multiple mRNA targets by a single sRNA. While we initially sought to determine kinetic parameters of regulation at translation and degradation levels for an sRNA that was classically categorized as a post-transcriptional regulator, we unexpectedly found that SgrS can regulate co-transcriptionally. Similar co-transcriptional regulation was also observed for the sRNA RyhB. Previous models of sRNA regulation were able to reproduce mRNA repression assuming only post-transcriptional regulation (*23*,*24*,*62*–*68*). The different order of sRNA and mRNA induction may explain why co-transcriptional regulation has not been noted in previous studies. Previous studies mostly either induced sRNAs in the presence of pre-existing or pre-induced mRNAs, or co-induced mRNAs and sRNAs simultaneously, whereas we pre-induced sRNAs to a certain level before inducing and tracking the changes of targets. Therefore, we created a time window, *i.e.*, early induction phase, when the mature mRNA level was low and the ratio between the nascent mRNA and the mature mRNA was high. Given the high abundance of pre-induced sRNA, and assuming in our model that sRNA used the same binding kinetics for both nascent and mature mRNA targets, the action of sRNA at the co-transcriptional level was enhanced compared to the cases where mature mRNAs were predominant. The effect of co-transcriptional regulation may be further enhanced by the target being plasmid-encoded. Because total target mRNA transcription was contributed by multiple plasmids in our experimental setting, the sRNA may more effectively regulate mRNA co-transcriptionally by targeting multiple transcription sites.

Although our target reporter mRNA genes are encoded by plasmids, it is very likely that sRNAs can act co-transcriptionally on the chromosomally encoded, endogenous genes. In order to co-transcriptionally regulate chromosomally encoded targets, sRNAs should be able to diffuse into the nucleoid region, which normally has a higher diffusion barrier. A previous report demonstrated that sRNAs have unbiased distribution between the nucleoid and cytoplasm using a few plasmid-encoded sRNAs as examples (*69*,*70*). Here, using single-molecule localization microscopy (SMLM), we confirmed the unbiased localization for these two sRNAs under our experimental conditions (Figure 7). In addition, the chaperone protein, Hfq, was observed to diffuse freely into the nucleoid region using single-particle tracking (*71*,*72*) and to bind to the nascent transcripts in a recent study using Chip-seq (*73*). It is likely that at least part of the Hfq binding to the nascent transcripts is mediated by sRNAs.

**Figure 7.**
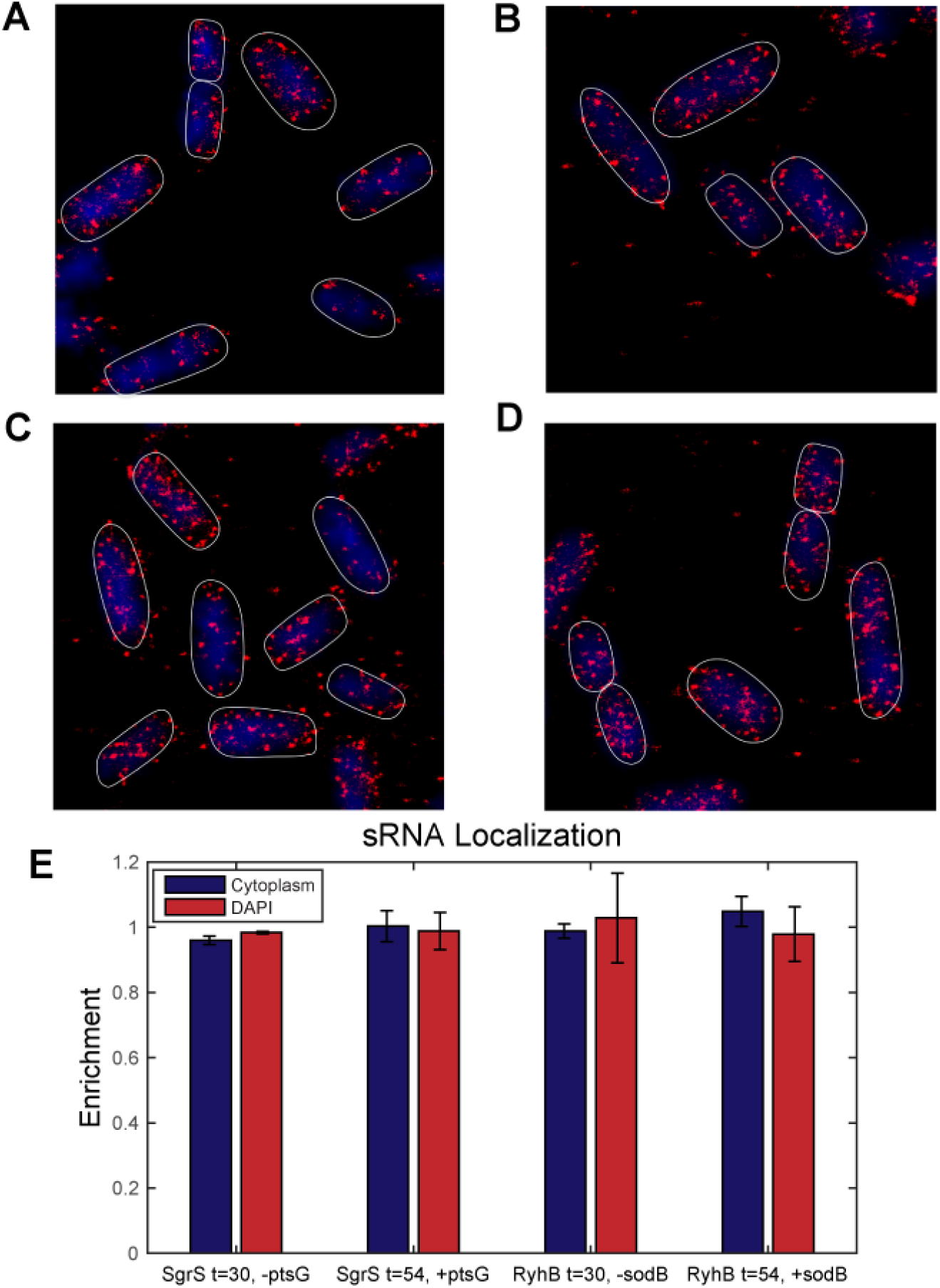
sRNA has unbiased localization between nucleoid and cytoplasm of bacterial cells. Representative SMLM images of SgrS (A) in the absence of *ptsG-sfGFP* mRNA induction (30 minutes after sRNA induction, before mRNA induced), (B) in the presence *ptsG-sfGFP* mRNA induction (54 minutes after sRNA induction, 24 minutes after mRNA induction). Representative SMLM images of RyhB (C) in the absence and (D) presence of *sodB*_130_*-sfGFP* induction. Red points are labeled sRNA detected by SMLM imaging. Blue region represents DAPI-stained nucleoid region. (E) The 3-dimensional localization for SgrS and RyhB in the absence and presence of the target mRNA was determined. Both SgrS and RyhB exhibited unbiased localization between the nucleoid and cytoplasm regardless of the presence of their target mRNAs.

While the majority of sRNAs are categorized as post-transcriptional regulators, cases have also been reported in which sRNAs can regulate transcription elongation, for example, by modulating the accessibility of the binding site of Rho factor, or by the conformational switch between terminator and antiterminator structures (*74*–*77*). Interestingly, previously characterized post-transcriptional sRNA regulators, DsrA, ArcZ and RprA can also regulate the target *rpoS* mRNA by suppressing pre-mature Rho-dependent transcription termination, a mechanism that may be widespread in bacterial genes with long 5’ UTR containing Rho binding site (*54*). In the case of SgrS and RyhB, there are no predicted Rho-independent termination sequences in the reporter mRNAs (*78*). However, we cannot exclude possibility of the presence of the Rho binding site. In addition, we observe that, for the same sRNA-mRNA pair, efficiency of co-transcriptional regulation is in general higher in the WT compared to the *rne*701 background (smaller *P* value in WT compared to *rne*701 background), indicative of a positive role for RNase E, or active degradation, in this process. However, we did not observe a positive correlation between *P* and *β*_e_, when comparing among different sRNA-mRNA pairs. We suspect that other unknown mechanisms may also contribute to the efficiency of co-transcriptional regulation. One possibility may be that formation of an sRNA-mRNA duplex at the exit tunnel of RNAP, or association of Hfq may inhibit transcriptional elongation. Nevertheless, our model suggests that sRNAs can act on nascent transcripts as soon as the sRNA binding sites are released from the RNA polymerases (Figure 8A).

**Figure 8.**
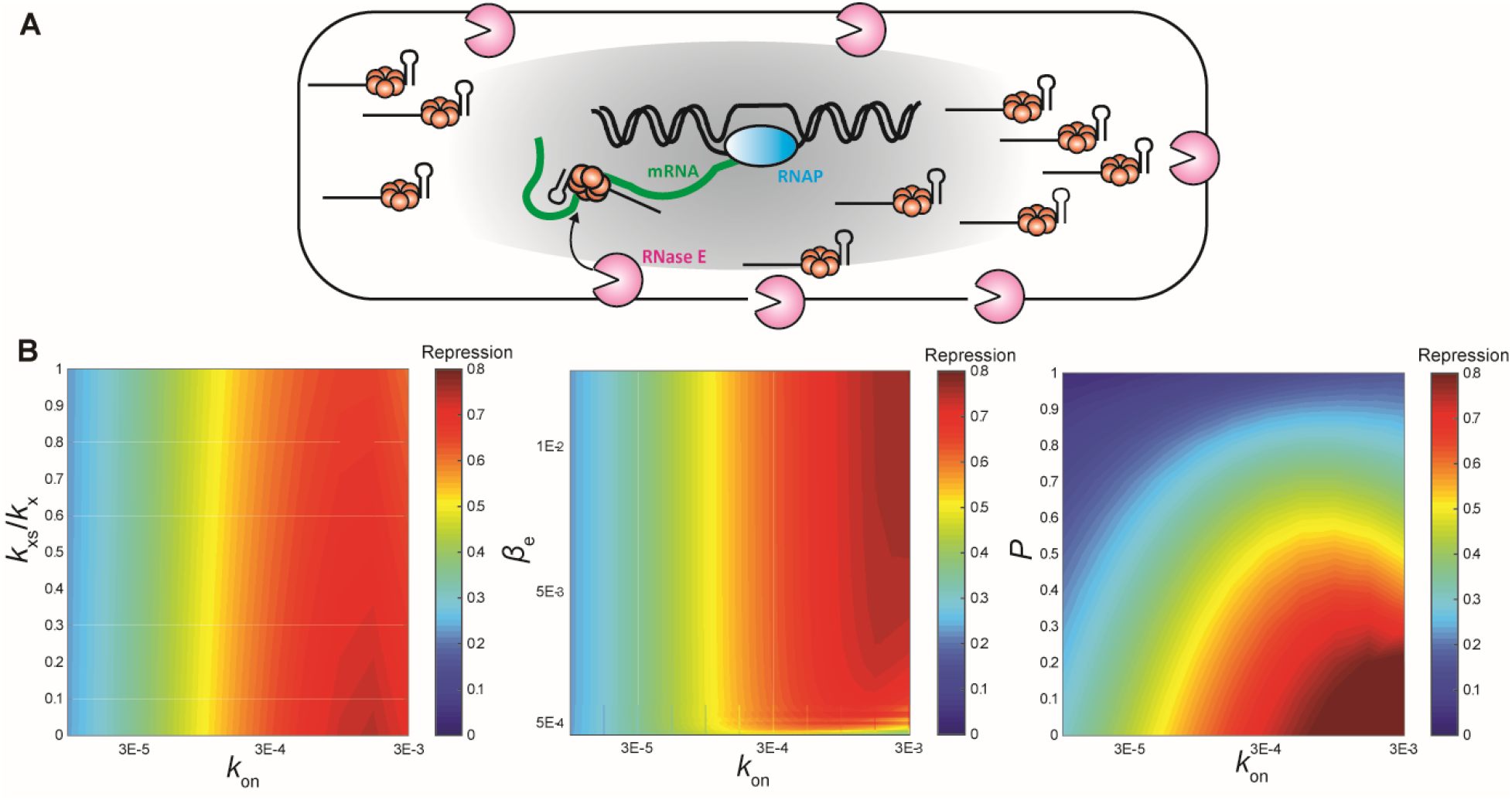
Model for co-transcriptional regulation by sRNAs. (A) sRNAs can freely diffuse in into the nucleoid region of bacterial cells, and bind to the target mRNAs as soon as the sRNA binding site is transcribed. Active degradation through recruitment of RNase E positively contributes to the efficiency of co-transcriptional regulation. (B) Protein-level repression heatmap, calculated by screening across the listed parameters. Repression level of 1 represents complete repression of protein expression; 0 means no repression. For the left panel, *β*_e_ = 1.0×10^−3^ and *P* = 0.32. For the middle panel, *k*_xs_/*k*_x_ = 0.5 and *P* = 0.32. For the right panel, *k*_xs_/*k*_x_ = 0.5 and *β*_e_ = 1.0×10^−3^. For all simulations, *k*_init_, *k*_x_, *k*_off_, *β*_m_, *β*_ms_, *β*_s_, and *k*_s_ were set to the measured or MAP values for *ptsG* (Table 1 and Supplementary Table S3).

Finally, our model suggests several kinetic steps that can determine the overall regulation efficiency. The binding kinetics between the sRNA and mRNA is the primary determinant of regulation efficiency. While *k*_off_ differs largely between different sRNAs, within the same regulon of a sRNA, *k*_on_ changes more dramatically compared to *k*_off_, and contributes to the regulation priority of different mRNAs by the same sRNA. At a constant *k*_on_, the strength of translation-level regulation (*k*_xs_/*k*_x_), sRNA-induced RNase E-mediated active degradation (*β*_e_), and regulation efficiency at co-transcriptional level (*P*) all positively contribute to the overall regulation efficiency. However, a fast *k*_on_ (>10^5^ M^−1^S^−1^) is generally needed to repress the target by more than 50% regardless of the rates or efficiencies at other steps (Figure 8B), suggesting that binding of the sRNA to the target mRNA might be the rate-limiting step. This is consistent with the observation that *purR*, which has a very low *k*_on_ rate, has the lowest regulation efficiency among SgrS regulon despite a higher *β*_e_. The binding kinetics are not correlated with the *in vitro* predicted hybridization thermodynamics (Supplementary Figure S1) *(79),* suggesting that more factors *in vivo* can affect the sRNA target search process. Interestingly, when comparing different sRNA-mRNA pairs, we found a positive correlation between *k*_on_ and the basal translation rate of the mRNA (*k*_x_) (Supplementary Figure S10), as noted previously (*63*). Specifically, higher *k*_on_ observed for *sodB*_130+30_ compared to *sodB*_130_ is possibly due to its higher *k*_x_. This correlation implies a potential positive role of translating ribosomes in promoting sRNA binding, perhaps through unwinding the secondary structures at the sRNA binding site (*80*). From a functional point-of-view, it is also logical to have a higher regulation efficiency on the most translated targets under stress conditions to achieve the most effective response.

## Data Availability

All data is deposited at dryad.org. All code for simulations and analysis is available at: https://github.com/JingyiFeiLab/Regulation_Kinetics

## Acknowledgements

We thank Dr. M.S. Azam for useful discussion, and MRSEC Shared User Facilities at the University of Chicago for 3D-printing service (NSF DMR-1420709 and DMR-2011854). We also thank the University of Chicago-operated Midway, a high performance computing cluster that forms the core of the Research Computing Center’s (RCC) computational infrastructure.

## Author contributions

M.A.R., C.K.V. and J.F. conceived the project and M.A.R. and J.F. designed the experiments. M.A.R., E.M.H., X.M. and M.B. conducted the experiments. M.A.R. and S.C. designed the models and performed the simulations with guidance from L.H. and A.R.D. M.A.R., C.K.V., and J.F. interpreted the data, M.A.R. and J.F. wrote the manuscript. All authors edited and approved the manuscript.

