## Supplementary information for "Kinetic model of small RNA-mediated regulation suggests that a small RNA can regulate co-transcriptionally"

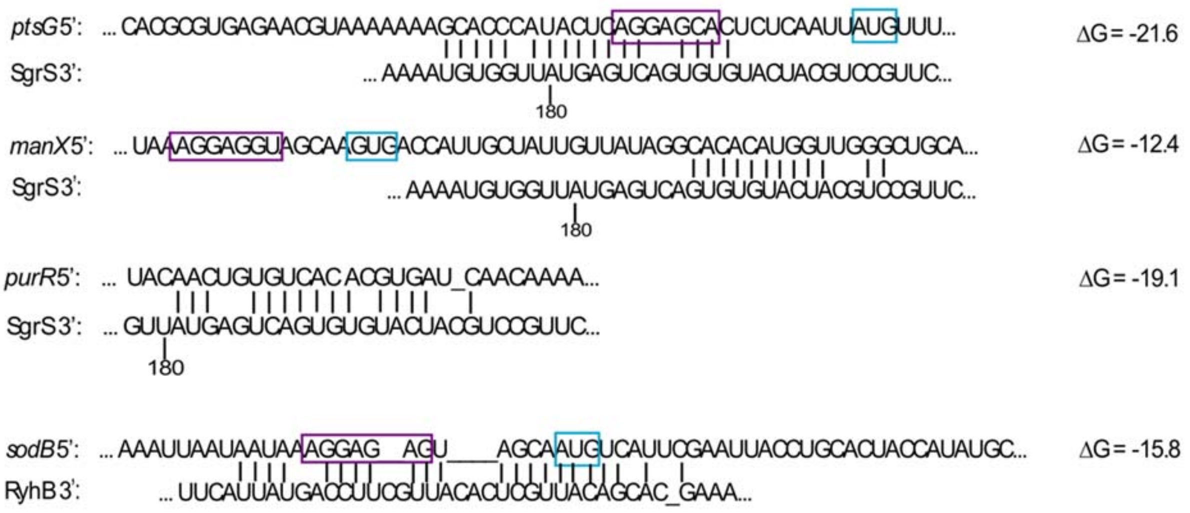

**Figure S1. sRNA target sequences.** Sequences of SgrS mRNA targets (*ptsG, manX*, and *purR*) and RyhB mRNA target (*sodB*). Vertical bars represent base pairing between the mRNA and sRNA. Purple boxes represent the Shine-Dalgarno sequence of the mRNA. Blue boxes represent the start codon of the mRNA-*sfGFP* fusion. The *purR* Shine-Dalgarno sequence and start codon are far upstream of the sRNA binding site. ΔG for each base pair interaction was calculated using the DINAMelt server (1)

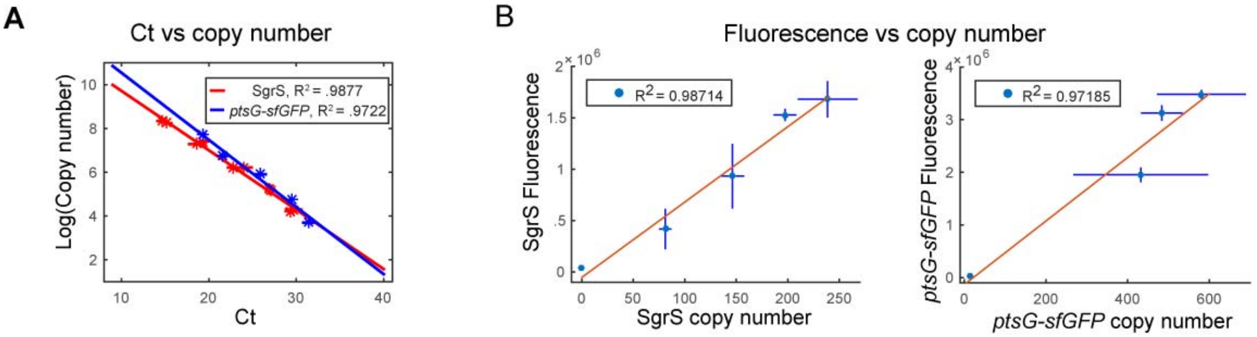

**Figure S2. RT-qPCR calibration for RNA copy number determination.** (A) Calibration curve relating copy number of the input RNA to the Ct value. Dilution series of *in vitro* transcribed SgrS and *ptsG-sfGFP* mRNA were spiked into the cell pellet of the Δ*sgrS* strain without the plasmid encoding the target mRNA. The spike-in RNAs and the cell pellet were put through total RNA extraction, RT and qPCR reactions and Ct values were determined. (B) Calibration curves relating the fluorescence value of each RNA from image to RNA copy number determined by RT-qPCR and calibration curve from (A). Error bars represent standard deviation of duplicate measurements.

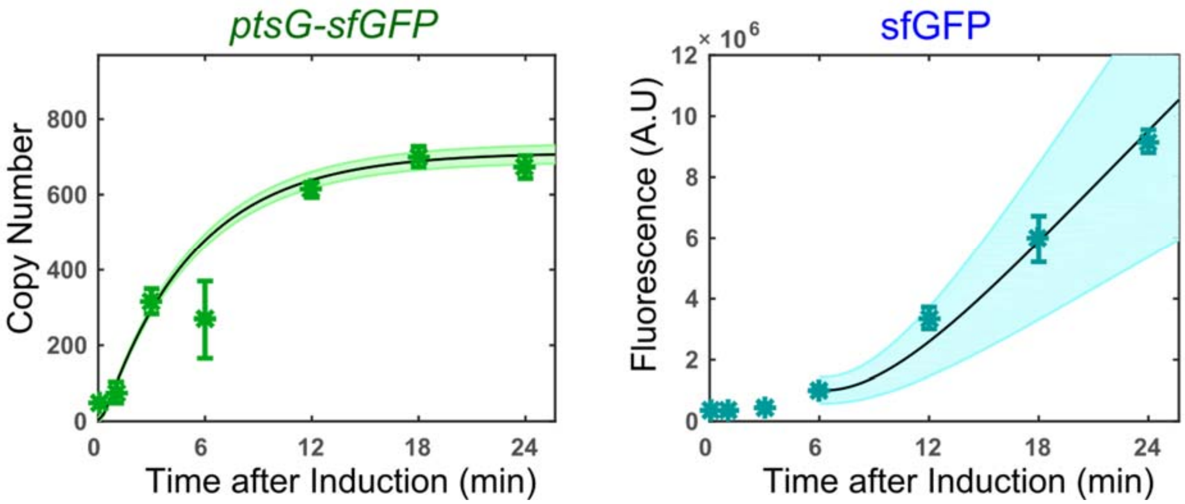

**Figure S3. Transcription and translation of *ptsG-sfGFP* in the absence of αMG in the WT strain.** The mRNA and protein levels are significantly higher than those in the Δ*sgrS* strain in the presence of αMG experiments (Figure 2C). Error bars represent standard deviation of duplicate measurements.

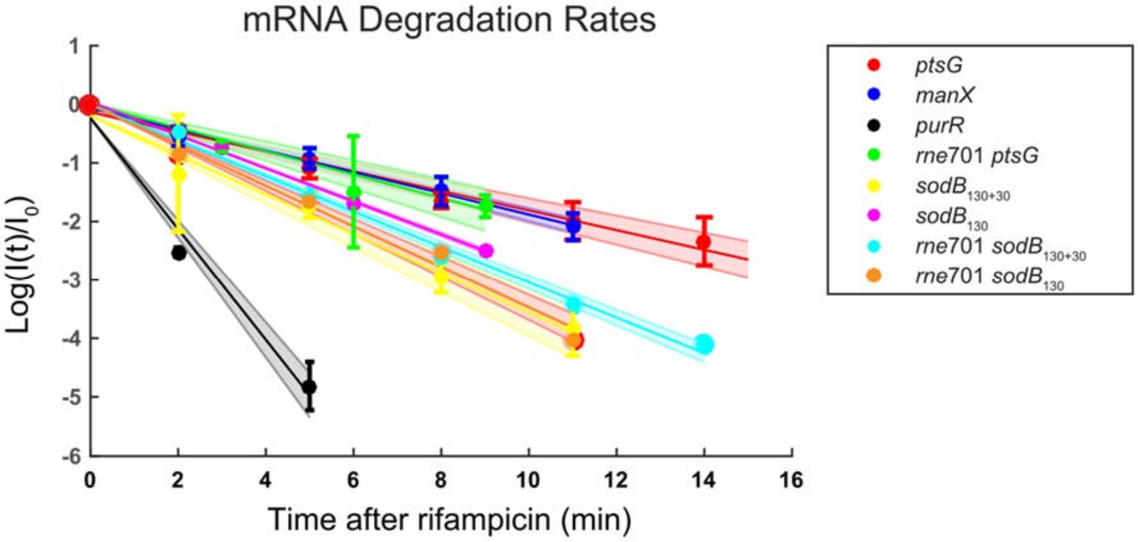

**Figure S4. Measurements of mRNA degradation rates.** Degradation rates for all target mRNAs used in this study were determined with rifampicin pulse chase experiments. Degradation rates were measured in both WT *rne* and *rne*701 backgrounds for *ptsG* and *sodB* constructs, confirming that the degradation rates for the target mRNA fusions were the same in both backgrounds. Y-axis shows the fluorescent intensity of the target mRNA relative to its intensity when rifampicin was added. Degradation rates were fit to an exponential decay model. Error bars represent standard deviation of duplicate measurements.

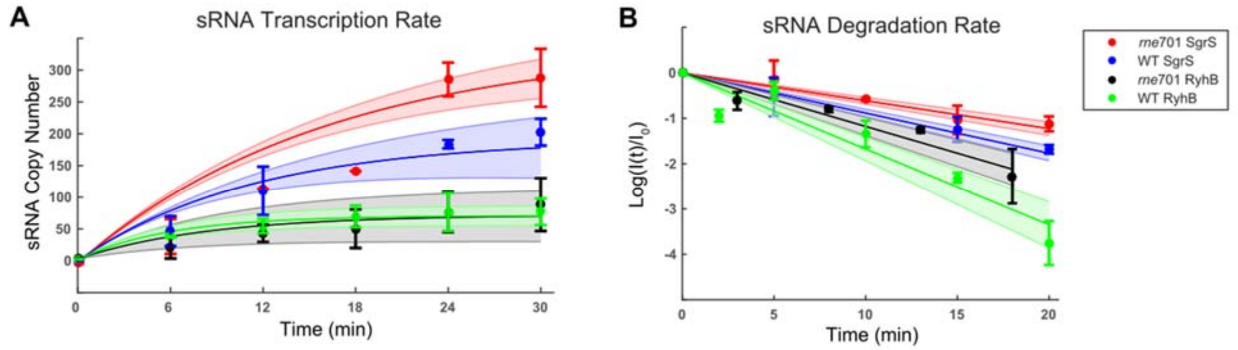

**Figure S5. Measurements of sRNA transcription and degradation rates.** (A) SgrS and RyhB transcription rates were measured in both WT *rne* and *rne*701 backgrounds by inducing the appropriate stress in the absence of target mRNA. (B) SgrS and RyhB degradation rates were measured in both WT *rne* and *rne*701 backgrounds by inducing sRNA to steady state levels, then removing the inducer from the media. Degradation rates were fit to an exponential decay model. Error bars represent standard deviation of duplicate measurements.

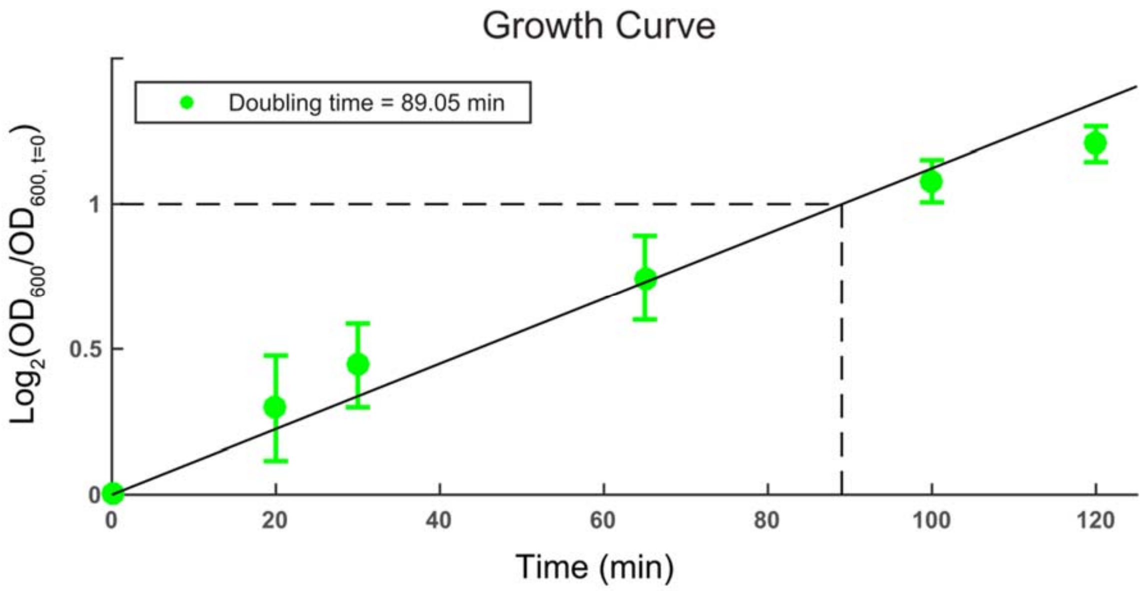

**Figure S6. Cell doubling time measurement.** $Log_2(OD_{600})$ relative to $OD_{600}$ at first time point in the exponential growth phase is plotted over time to determine cell doubling time. Error bars represent standard deviation of duplicate measurements.

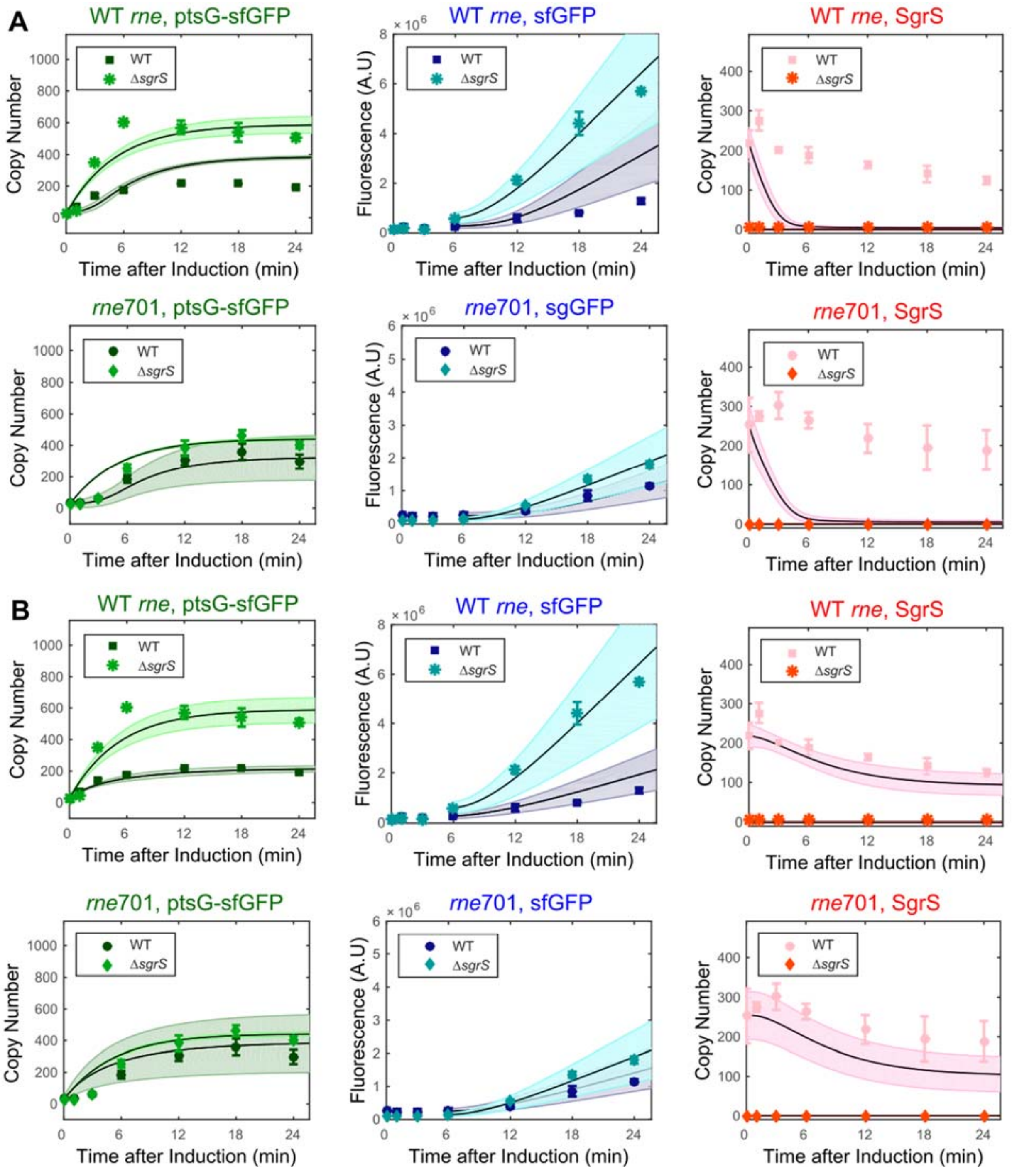

**Figure S7. Fits of SgrS regulation on *ptsG-sfGFP* using one-step transcription module.** (A) Best fit of post-transcriptional regulation model for *ptsG*. (B) Best fit of uncoupled co-transcriptional regulation model for *ptsG*.

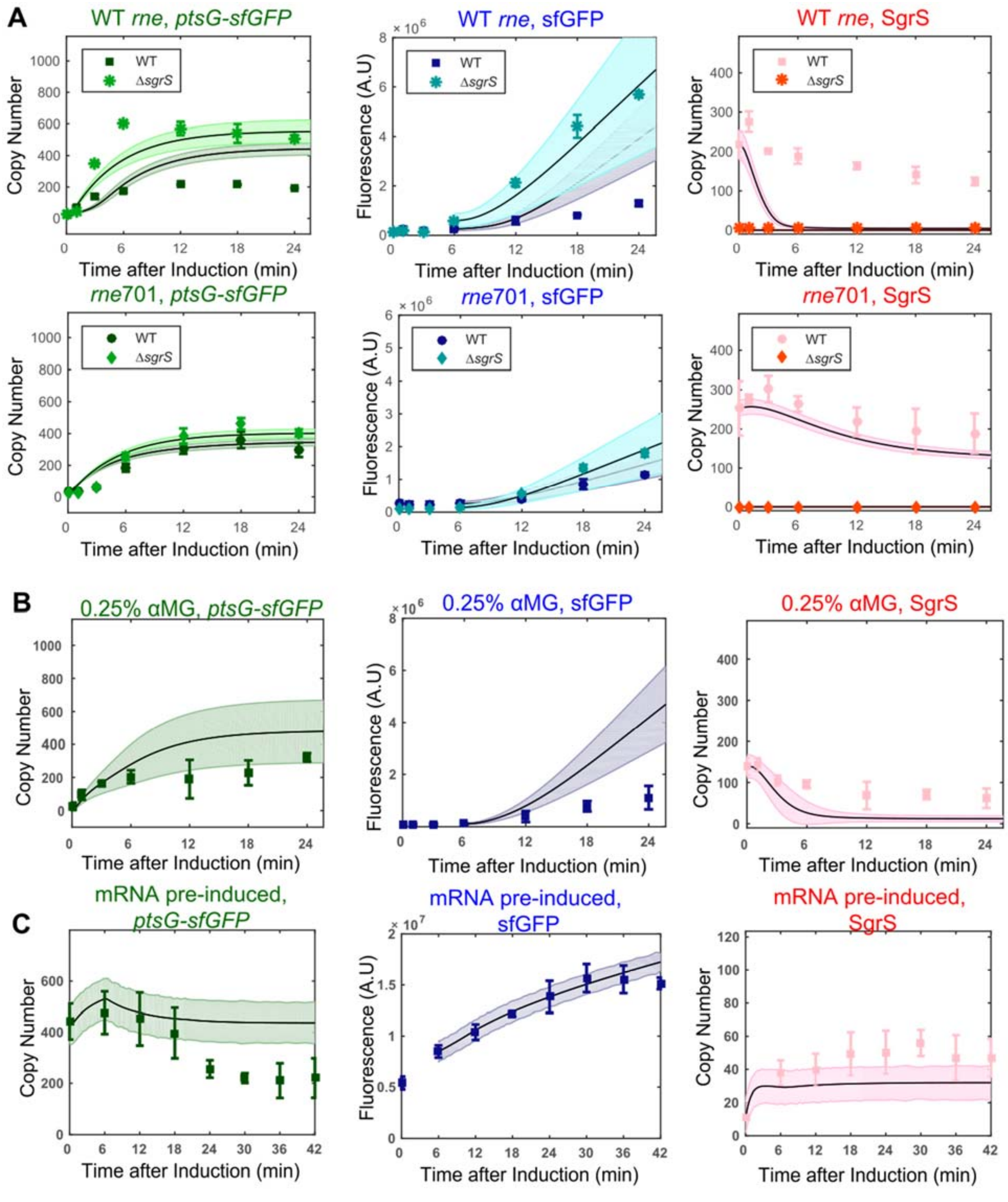

**Figure S8. Fits and predictions of post-transcriptional regulation model for SgrS regulation on *ptsG-sfGFP* using two-step transcription module.** (A) Best fit of post-transcriptional regulation model *ptsG* in WT and *rne701* background. (B) Simulation of best fit using post-transcriptional model on 0. 25% αMG validation dataset. (C) Simulation of best fit using post-transcriptional model on pre-induced mRNA dataset.

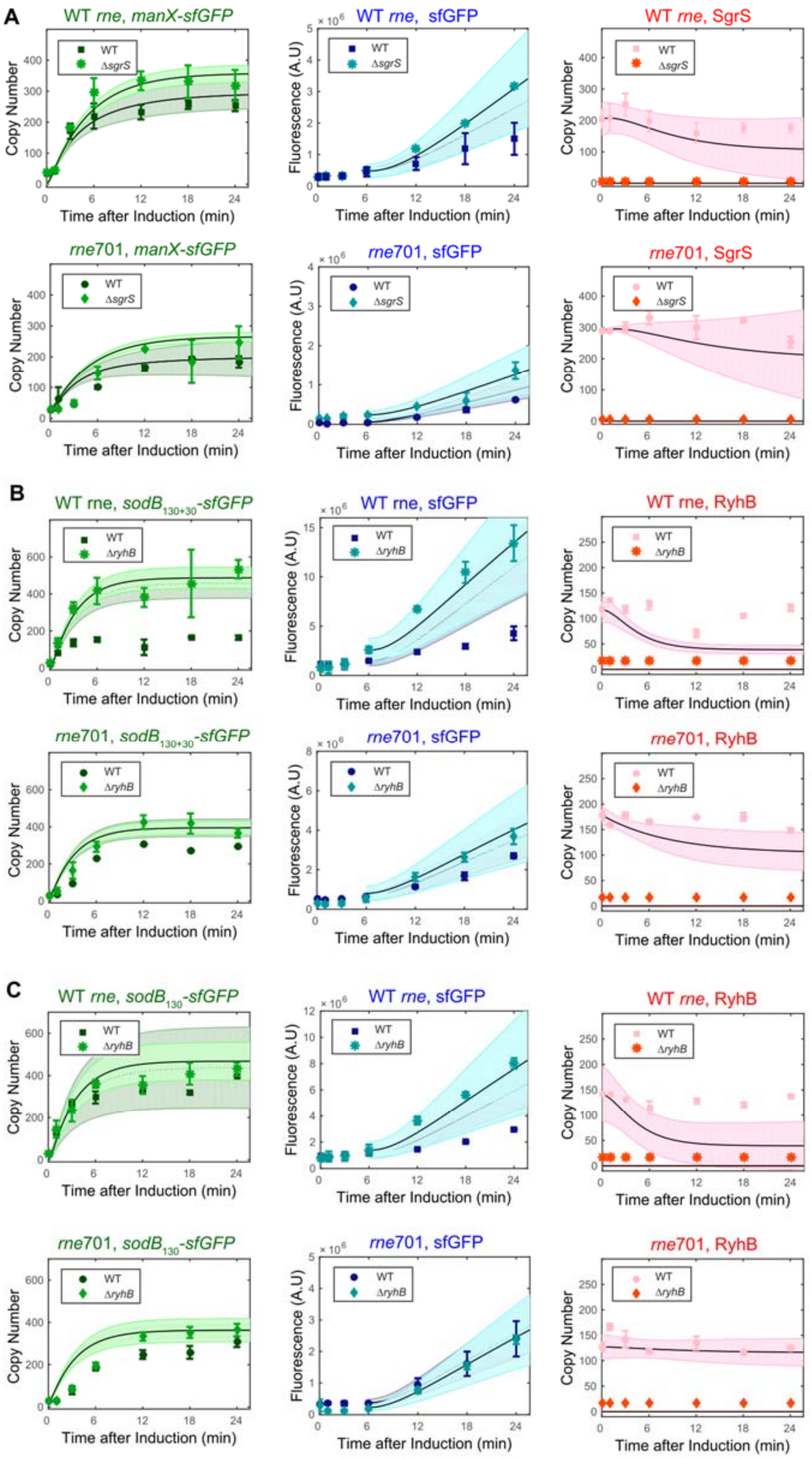

**Figure S9. Post-transcriptional model fits for other sRNA-mRNA pairs.** (A) Best fit of post-transcriptional regulation model in for SgrS regulation of *manX-sfGFP.* (B) Best fit of post-transcriptional regulation model in for RyhB regulation of *sodB*$_{130+30}$*-sfGFP* (C) Best fit of post-transcriptional regulation model in for RyhB regulation of *sodB*$_{130}$*-sfGFP.*

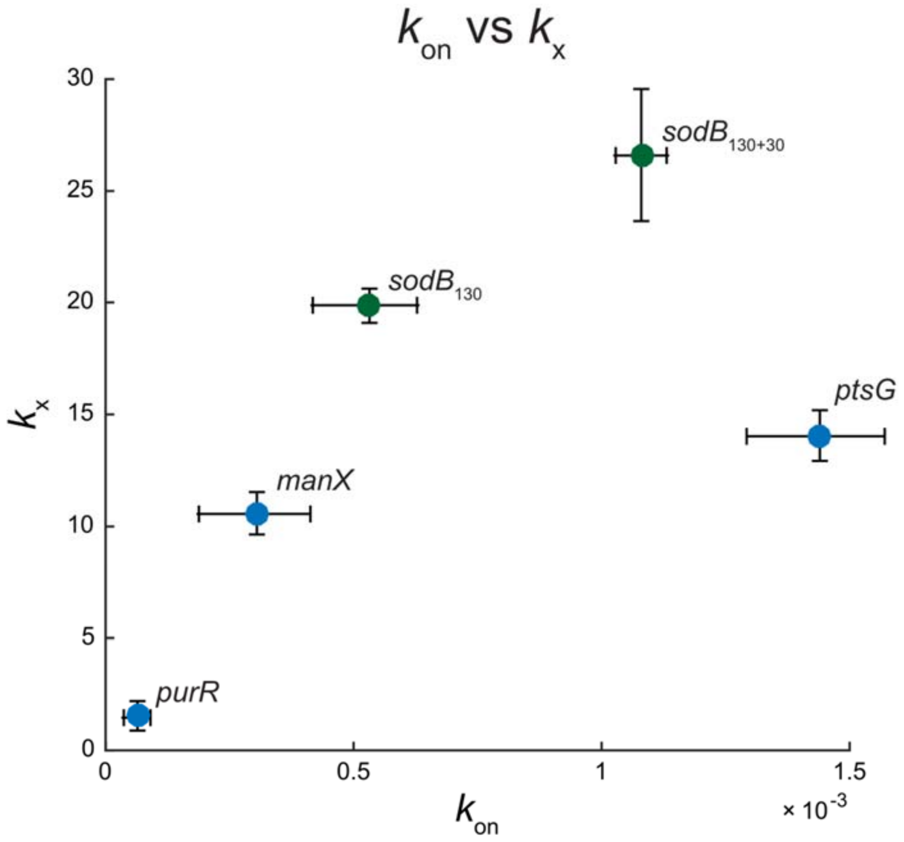

**Figure S10. Translation rate vs on-rate of sRNA binding.** $k_x$ *vs.* $k_{on}$ for all mRNA targets. Points in blue are mRNA targets of SgrS. Points in green are targets of RyhB. Error bars represent standard deviation of calculated MAP values (see Table 1 and Supplementary Table S3).

**Table S1. List of all strains and plasmids used in this study**

| Strain | Description | Reference |
|---|---|---|
| DB166 | *WTsgrS, λattB::lacIq,tetR, specR* | This study |
| JH111 | *ΔsgrS, λattB::lacIq,tetR, specR* | Reference (2) |
| XM100 | *lacIq tetR specR rne701-FLAG::cat* | This study |
| XM101 | *ΔsgrS lacIq tetR specR rne701-FLAG::cat* | This study |
| XM221 | *lacIq, tetR, specR, rne701-FLAG::cat, ryhB::tet* | This study |
| DB186 | *lacIq, tetR, specR, ryhB::cat* | This study |
| **Plasmids** | **Description** | **Reference** |
| | pSMART_*ptsG*-10aa-*sfGFP* | This study; modified from Reference (3) |
| | pSMART_*manX*-34aa-*sfGFP* | This study; modified from Reference (4) |
| | pSMART_*purR*-32aa-*sfGFP* | This study |
| | pSMART_*sodB*430-sfGFP | This study |
| | pSMART_*sodB*130-sfGFP | This study |
| | pSMART_*sodB*130+30-sfGFP | This study; modified from Reference (5) |

### Table S2. List of all oligonucleotides used in this study

| Strain/plasmid generation primers | Sequence 5'-3' | Description |
|---|---|---|
| JZ25 | ATCCCTATCAGTGATAGAGATACTGGAGCACAGAATTCATAAATAAAGGG | Tet promoter + ptsG |
| JZ26 | GTCAATCTCTATCACTGATAGGGACTTTCTCGAGGTGAAGACGAAA | Tet promoter + pZEMB vector |
| EH1 | TCGTCTTCACCTCGAGAAAGTC | Amplifies tet_mRNA-sfGFP for ligation into pSMART |
| EH2 | CGAACGCCTAGGTCTAGGGCGG | Amplifies tet_mRNA-sfGFP for ligation into pSMART |
| EH3 | ATCCCTATCAGTGATAGAGATACTGGATACTGAGCACAGAATTC | Tet promoter + manX |
| EH307 | CTGGACCTGGGGATCCGCTGGCTCCG | sodB430 + pSMART |
| EH308 | CCAGCGGATCCCCAGGTCCAGCCAGAACCAAAG | pSMART + sodB430 |
| EH309 | TATTGTGCGTATGAATTCTGTGCTCCAGTATCTCTATCACTG | sodB + pSMART |
| EH310 | AGCACAGAATTCATACGCACAATAAGGCTATTGTACGTATG | pSMART + sodB |
| EH390 | CTCGATGGTTTCCGCAGAAATGTG | sodB130 |
| EH391 | GGATCCGCTGGCTCCGC | sodB130 |
| EH440 | ATCAAAAACTTTGGTTCTGGCTGG | sodB130+30 |
| EH441 | CTCGATGGTTTCCGCAGAAA | sodB130+30 |
| OXM211 | GTGTTGGACAAGTGCGAATGAGAATGATTATTATTGTCTCCATTAATTCCTAATTTTTGTTGACACTCTATC | |
| OXM212 | AAGCACTCCCGTGGATAAATTGAGAACGAAAGATCAAAAAGAATAACATCATTTGGTGACGAAATAACTA | |
| OXM112 | ATGAGCAAAGGAGAAGAAC | pSMART |
| OXM113 | GAATTCTGTGCTCCAGTATC | pSMART |
| OXM115 | agttcttctcctttgctcatGAATTCGCCAGAACCAGC | purR + pSMART |
| OXM116 | gatactggagcacagaattcTACACTATTTGCGTACTGGC | purR + pSMART |
| **qPCR primers** | **Sequence 5'-3'** | |
| ptsG_U_F | CAGAATTCATAAATAAAGGGCGCTTAGA | qPCR targeting *ptsG-sfGFP* upstream of SgrS binding site |
| ptsG_U_R | TCTCACGCGTGGCAAGG | qPCR/RT targeting *ptsG-sfGFP* upstream of SgrS binding site |
| ptsG_D_F | CCGTTCAACTAGCAGACCATTA | qPCR targeting *ptsG-sfGFP* downstream of SgrS binding site |
| ptsG_D_R | GACAGATTGTGTCGACAGGTAA | qPCR/RT targeting *ptsG-sfGFP* downstream of SgrS binding site |
| 16S rRNA_F | AGGCCTTCGGGTTGTAAAGT | qPCR targeting ribosomal RNA |
| 16S rRNA_R | ATTCCGATTAACGCTTGCAC | qPCR/RT targeting ribosomal RNA |
| SgrS_F | AGCGTCCCACAACGATTAAC | qPCR targeting SgrS |
| SgrS_R | CACCAATACTCAGTCACACATGA | qPCR targeting SgrS |
| **In vitro transcription primers** | **Sequence 5'-3'** | |
| SgrS +T7_F | TAATACGACTCACTATAGGGATGAAGCAAGGGGGTGC | |
| SgrS +T7_R | AAAAAAAACCAGCAGGTATAATCTGCT | |

| | |
|---|---|
| ptsG-sfGFP +T7 F | TAATACGACTCACTATAGGCACAGAATTCATAAATAAAGGGCG |
| ptsG-sfGFP +T7 R | CCGCCCTAGACCTAGGCGTTCG |
| **FISH Probes** | **Sequence 5'-3'** |
| SgrS_1 | GTGCTGATAAAACTGACGCA |
| SgrS_2 | ACTTCGCTGTCGCGGTAAAA |
| SgrS_3 | CTTAACCAACGCAACCAGCA |
| SgrS_4 | CATGGTTAATCGTTGTGGGA |
| SgrS_5 | ATCCCACTGCATCAGTCCTT |
| SgrS_6 | GTCAACTTTCAGAATTGCGG |
| SgrS_7 | TCAGTCACACATGATGCAGG |
| SgrS_8 | GCGGGTGATTTTACACCAAT |
| SgrS_9 | AACCAGCAGGTATAATCTGC |
| sfGFP_1 | ATTTGTGCCCATTAACATCA |
| sfGFP_2 | GAGTAGTGACAAGTGTTGGC |
| sfGFP_3 | TCATGTGATCCGGATAACGG |
| sfGFP_4 | TAGTGCGTTCCTGTACATAA |
| sfGFP_5 | GCCGTGATGTATACATTGTG |
| sfGFP_6 | GTTAGCTTTGATTCCATTCT |
| sfGFP_7 | GCTAGTTGAACGGAACCATC |
| sfGFP_8 | CGCCAATTGGAGTATTTTGT |
| sfGFP_9 | TGTCGACAGGTAATGGTTGT |
| sfGFP_10 | TCAAGAAGGACCATGTGGTC |
| RyhB_1 | GCGAGGGTCTTCCTGATCGC |
| RyhB_2 | ATGTCGTGCTTTCAGGTTCT |
| RyhB_3 | AATACTGGAAGCAATGTGAG |
| RyhB_4 | GCCAGCACCCGGCTGGCTAA |
| 16S rRNA | CCC CAG TCA TGA ATC ACA AA |

**Table S3. Additional kinetic parameters**

| | | | |
|---|---|---|---|
| **SgrS** | | | |
| $\alpha_s$ **(molecule•s$^{-1}$)** | 0.36 ± 0.03 | | |
| $\beta_s$ (WT) **(s$^{-1}$)** | (1.5 ± 0.3) x$10^{-3}$ | | |
| $\beta_s$ (*rne*701) **(s$^{-1}$)** | (1.0 ± 0.1) x$10^{-3}$ | | |
| **Targets of SgrS** | ***ptsG*** | ***manX*** | ***purR*** |
| $\alpha_m$ (Δ*sgrS* + αMG) **(molecule•s$^{-1}$)** | 0.09 ± 0.01 | 0.06 ± 0.01 | 0.08 ± 0.01 |
| $\alpha_m$ (*rne*701Δ*sgrS* + αMG) **(molecule•s$^{-1}$)** | 0.07 ± 0.01 | 0.04 ± 0.01 | 0.08 ± 0.01 |
| $k_x$ (Δ*sgrS*) **(AU•s$^{-1}$)** | 14 ± 1 | 11 ± 1 | 1.5 ± 0.7 |
| $k_x$ (*rne*701Δ*sgrS*) **(AU•s$^{-1}$)** | 6.1 ± 0.4 | 5.46 ± 0.03 | 0.7 ± 0.3 |
| $\beta_m$ **(s$^{-1}$)** | (3.2 ± 0.1) x$10^{-3}$ | (3.3 ± 0.1) x$10^{-3}$ | (1.7 ± 0.1) x$10^{-2}$ |
| **RyhB** | | | |
| $\alpha_s$ **(molecule•s$^{-1}$)** | 0.26 ± 0.07 | | |
| $\beta_s$ (WT) **(s$^{-1}$)** | (2.8 ± 0.2) x$10^{-3}$ | | |
| $\beta_s$ (*rne*701) **(s$^{-1}$)** | (2.0 ± 0.4) x$10^{-3}$ | | |
| **Targets of RyhB** | ***sodB$_{130+30}$*** | ***sodB$_{130}$*** | |
| $\alpha_m$ (Δ*ryhB* + DIP) **(molecule•s$^{-1}$)** | 0.11 ± 0.03 | 0.12 ± 0.01 | |
| $\alpha_m$ (*rne*701Δ*ryhB* + DIP) **(molecule•s$^{-1}$)** | 0.11 ± 0.01 | 0.09 ± 0.02 | |
| $k_x$ (Δ*ryhB*) **(AU•s$^{-1}$)** | 27 ± 2 | 19.9 ± 0.9 | |
| $k_x$ (*rne*701Δ*ryhB*) **(AU•s$^{-1}$)** | 10 ± 2 | 8 ± 1 | |
| $\beta_m$ **(s$^{-1}$)** | (5.2± 0.1) x$10^{-3}$ | (5.0 ± 0.1) x$10^{-3}$ | |

**Table S4. Parameter prior distributions**

| | |
|---|---|
| **$\alpha_m$ (molecule•s$^{-1}$)** | $N(\mu_a,\sigma_a)$[a] |
| **$K_x$ (AU•s$^{-1}$)** | $N(\mu_k,\sigma_k)$ |
| **$k_{on}$ (molecule$^{-1}$s$^{-1}$)** | $U(10^{-9},10^{-2})$[b] |
| **$k_{off}$ (molecule•s$^{-1}$)** | $U(10^{-5},10)$ |
| **$k_{xs}/k_x$** | $U(0,1)$ |
| **$\beta_{ms}$ (s$^{-1}$)** | $U(\beta_m,1.0)$ |
| **$\beta_e$ (s$^{-1}$)** | $U(10^{-6},1.0)$ |

[a] Normal distribution with mean and standard deviation calculated from –sRNA experiments
[b] Uniform distribution
